# Experience dependent contextual codes in the hippocampus

**DOI:** 10.1101/864090

**Authors:** Mark H. Plitt, Lisa M. Giocomo

**Affiliations:** Department of Neurobiology, Stanford University, Stanford CA USA

## Abstract

The hippocampus is a medial temporal lobe brain structure that contains circuitry and neural representations capable of supporting declarative memory. Hippocampal place cells fire in one or few restricted spatial locations in a given environment. Between environmental contexts, place cell firing fields remap (turning on/off or moving to a new spatial location), providing a unique population-wide neural code for context specificity. However, the manner by which features associated with a given context combine to drive place cell remapping remains a matter of debate. Here we show that remapping of neural representations in region CA1 of the hippocampus is strongly driven by prior beliefs about the frequency of certain contexts, and that remapping is equivalent to an optimal estimate of the identity of the current context under that prior. This prior-driven remapping is learned early in training and remains robust to changes in behavioral task-demands. Furthermore, a simple associative learning mechanism is sufficient to reproduce these results. Our findings demonstrate that place cell remapping is a generalization of representing an animal’s location. Rather than simply representing location in physical space, the hippocampus represents an optimal estimate of location in a multi-dimensional stimulus space.

## Main Text

The neural firing of hippocampal place cells often strongly correlates with an animal’s current spatial location in an environment, providing a neural basis for the brain’s code for space^1^. Between environmental contexts, place cell firing fields can appear, disappear or move – phenomena collectively referred to as ‘remapping’^2–10^. This reorganization in the firing locations of place cells results in unique population-wide representations for different environmental contexts. However, the factors that drive remapping remain incompletely understood. Here, we consider the idea that the animal’s prior beliefs about the frequency with which it encounters stimuli determine remapping patterns^11^. We find that context specific spatial codes are activated in a way that allows an animal to optimally estimate its location in the multi-dimensional stimulus space that determines ‘context’. We further show that this result can be accounted for by simple and long-standing models of hippocampal associative memory. This work provides the first quantitative framework for making precise predictions of how hippocampal population codes are formed and recruited across contexts.

### Prior beliefs about stimulus frequency determine CA1 place cell remapping

To examine how prior experience affects remapping of hippocampal representations, we performed two-photon imaging of CA1 pyramidal cells as mice traversed a number of visually similar virtual reality (VR) linear tracks, which were presented with different frequencies. We designed the VR tracks such that three dominant visual features of the environment could gradually blend (i.e. morph) between two extremes (visual features = orientation and frequency of bars on the wall, background color, color of tower landmarks) ^3,6,10^. The degree of blending in each feature is the coefficient of the affine combination of two values of that stimulus, referred to as the ‘morph value’, *S* (Fig 1a, Methods). For each trial, a shared morph value, 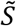, was chosen for all features from one of five values (0, .25, .5, .75, and 1) and then a jitter (uniformly distributed between -.1 to .1) was added to each feature independently. The overall morph value of a trial is the sum of the shared morph value and each of the jitters (*S* = -.3 to 1.3). Weber-Fechner’s law suggests that the animals will perceive log-fold rather than linear changes in the stimuli, so we considered the “log-morph” of a trial for some analyses (see Methods)^12^. In all sessions, mice performed a random foraging task, which required them to lick for water at a reward cue that appeared at a random location on the back half of the track (Extended Data Fig 1 and <u>Supplementary Video 1</u>). In each session, we were able to image several hundred to thousands of putative CA1 pyramidal cells simultaneously (98 – 2,149 cells simultaneously recorded; Fig 1b and Extended Data Fig 1f-g)

**Figure 1:**
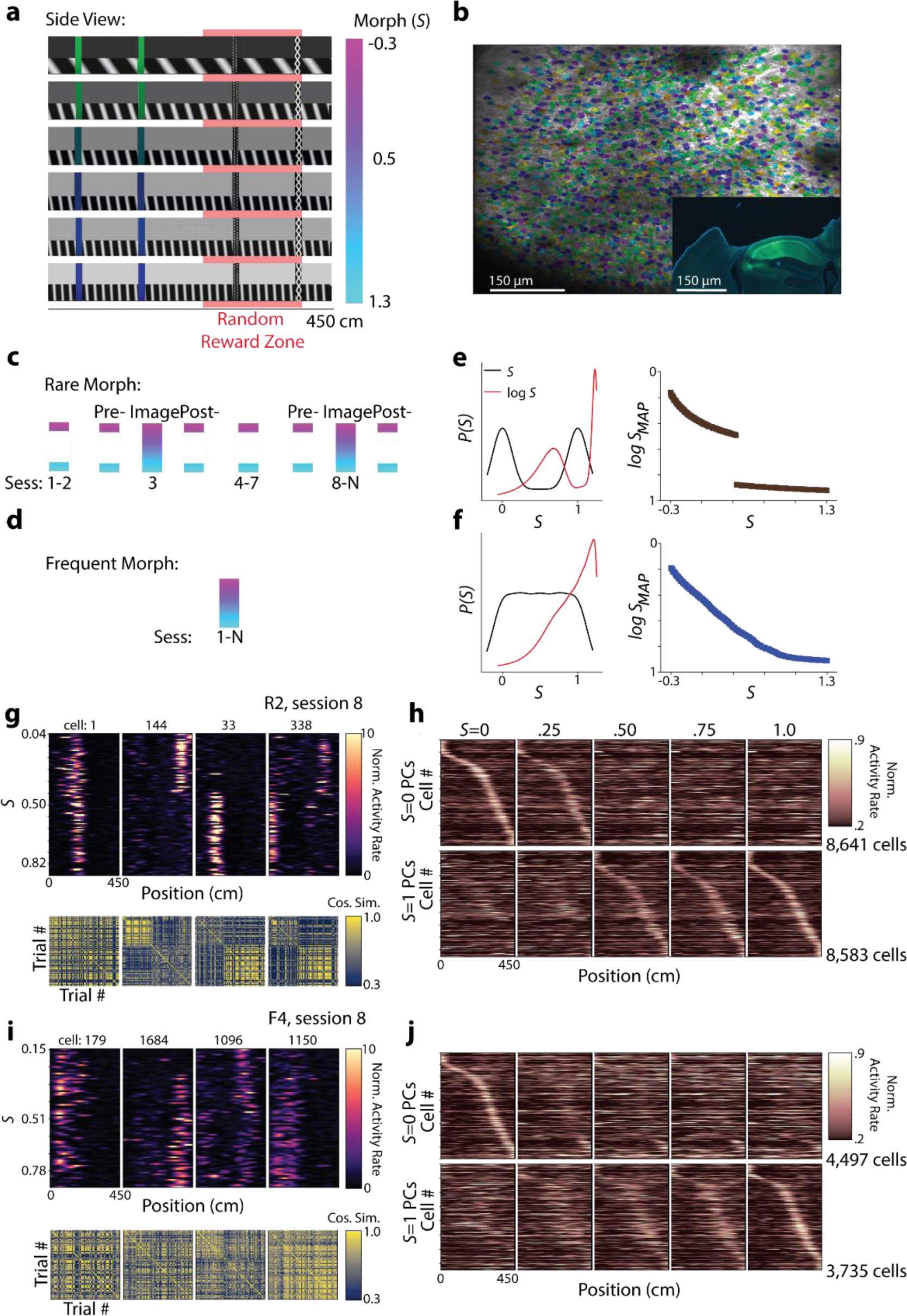
Training animals with different prior beliefs about stimulus frequency results in distinct CA1 remapping patterns. **a**, Stimulus design with example tracks for different morph (*S*) values. Side view of the track is shown. The color bar (right) indicates where each track falls along the morph axis, which is determined by the color of the background (dark to light grey), the color of the first two towers (green to blue) and the orientation and frequency of the sine waves on the walls (low frequency tilted sine waves to high frequency vertical sine waves). Liquid rewards appeared at a random location between 250 and 400 cm along the virtual track (red shaded region). **b**, Example mean motion-corrected image of the field of view from one imaging session (mouse F2, session 12; *λ*=920 nm) with identified CA1 neurons highlighted in different colors (Suite2p, n = 2004 cells). Inset shows a coronal view of the histology from the same animal (Green = GCaMP; Blue = DAPI). **c**, Illustration of the training protocol for the Rare Morph condition (n = 4 animals, R1-R4), color coded for morph (*S*) value as in (**a**). Imaging was performed on sessions (Sess.) 1, 3, and 8-N (Extended Data Fig 1f). Intermediate morph values are only shown on sessions 3 and 8-N, during which imaging also occurred. **d**, Illustration of the training protocol for the Familiar Morph condition (n = 5 animals, F1-F5). Imaging was performed on sessions 1, 3, and 8-N (Extended Data Fig 1g). Intermediate morph values are shown on each session. **e**, Rare Morph condition. *Left:* The idealized prior distribution over the morph values (black) and log-corrected morph values (red) experienced by the Rare Morph animals, *P(S). Right:* The log-corrected *Maximum a Posteriori* estimate of the stimulus, *log SMAP*, as a function of the true stimulus (morph value), *S*. **f**, Same as (**e**) but for the Frequent Morph condition. **g**, *Top Row:* Co-recorded place cells from an example Rare Morph session (mouse R2, session 8). Each column corresponds to the heat map for a different cell. Each row in the heat map indicates the activity of that cell on a single trial as a function of the position of the mouse on the virtual track (n=120 trials). Color code indicates deconvolved activity rate normalized by the overall mean activity rate for the cell (color bar far right). Rows are sorted by increasing morph value. *Bottom row:* Trial by trial cosine similarity matrices for the corresponding cell above, color coded for maximum (yellow) and minimum (blue) values. **h**, *Top row:* For all Rare Morph sessions 8-N, place cells were identified in the *S* = 0 morph trials and sorted by their location of peak activity (leftmost panel, n = 8,641 cells). Each row indicates the activity rate of a single cell as a function of position, averaged over all 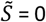 morph trials. Color indicates activity rate normalized by the peak rate from the 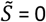 average activity rate map for that cell (color bar far right). The trial-averaged firing rate map for these cells is then plotted for binned morph values using the same sorting and normalization (remaining panels to the right). *Bottom Row:* For the same Rare Morph Sessions, place cells were identified in the 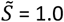 morph trials sorted by their location of peak activity (rightmost panel, n = 8,583 cells), and normalized by their peak value in the 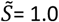 mean activity rate map. The same sorting and normalization is then used for the remaining panels to the left. **i**, Same as in (**g**) for an example Frequent Morph session (mouse F4, session 8, n = 75 trials). **j**, Same as in (**h**) for all Frequent Morph sessions 8-N (top row n = 4,497 cells; bottom row n = 3,735 cells).

We considered two training conditions. In the Rare Morph condition (n = 4 mice, R1 – R4), mice experienced very few trials with intermediate morph values (shared 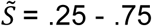) during the first seven training sessions (< 4% of trials with intermediate morph values before session 8; Fig 1c), giving these animals a bimodal prior over the (log-)morph values (idealized prior Fig 1c and empirical prior per mouse Extended Data Fig 2). By contrast, in the Frequent Morph condition (n = 5 mice, F1 – F5), mice were frequently exposed to trials with intermediate morph values (60% of trials with intermediate morph values before session 8; Fig 1e), giving these animals a flat prior over the morph values (except the tails of the distribution) and a ramping prior over log-morph values (idealized prior Fig 1f & empirical prior per mouse Extended Data Fig 2). Given these priors, an observer performing probabilistic inference would make different predictions about the identity of intermediate stimuli. In particular, the *Maximum a Posteriori* (MAP) estimate of the stimulus (i.e. choosing the most likely stimulus from the posterior distribution), is discontinuous in the Rare Morph condition (“Rare Morph MAP estimate curve”, Fig 1e) and continuous in the Frequent Morph condition (“Frequent Morph MAP estimate curve”, Fig 1f). MAP estimation in the Frequent Morph condition is also nearly identical to Maximum Likelihood estimation before the log morph correction.

We focused on CA1 remapping in sessions 8-N, as animals have had 7 sessions to develop a stable prior for the frequency of different stimuli (See Extended Data Fig 1 for number of sessions per mouse). In both the Frequent and Rare Morph condition, large numbers of place cells tiled the track with place fields (Rare = 13,985 place cells, 49.23% of recorded population; Frequent = 7,192 place cells, 34.02% of recorded population). For environments at the extreme morph values (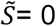 and 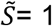), the proportion of cells with a place field in both environments slightly exceeded the proportion expected by randomly choosing cells with replacement from the population (11.4% and 4.92% of recorded cells with significant fields in both environments for Rare Morph and Frequent Morph sessions respectively; chance 9.12% and 3.76%, Extended Data Fig 3). This suggests that the place representations between the environments at the extreme morph values were largely independent in both the Frequent and Rare Morph conditions. Consistent with the probabilistic inference framework introduced above, the transition between these representations clearly differed between the Rare and Frequent Morph condition. In the Rare Morph condition, place cells maintained their representation until a threshold morph value (*S* = .25 – .50) and then coherently switched to a different representations (Fig 1g-h). In the Frequent Morph condition, place cells gradually changed their representation across intermediate morph values (Fig 1i-j).

### Discrete versus Continuous Representations of Context in Population Codes

We next sought to confirm that the remapping patterns we observed in place cells reflected the dominant patterns in the entire CA1 population in an unsupervised manner and included all identified cells in the subsequent analyses. To examine how single cell firing rate maps changed across morph values, we performed cross-validated non-negative matrix factorization (NMF)^13^ on the single cell trial × trial similarity matrices (Fig 2a). Intuitively, factors produced by the model can be thought of as ‘prototypical similarity matrices’. In the Rare Morph condition, a two factor NMF model yields one factor in which cells maintained a representation for a lower morph values and a second factor in which cells maintained a representation for a higher morph value (example session in Fig 2b, across sessions and mice in Fig 2c). On the other hand, a two factor NMF model for Frequent Morph condition yields factors in which cells show gradual changes in similarity across morph values (example session in Fig 2d, across sessions and mice in Fig 2e). This confirms that the place cell remapping observed in the average place cell population firing rate maps (Fig 1i-j) are also the dominant remapping patterns followed by all recorded hippocampal CA1 cells (see Extended Data Fig 4 for higher rank NMF models and model error analysis).

**Figure 2:**
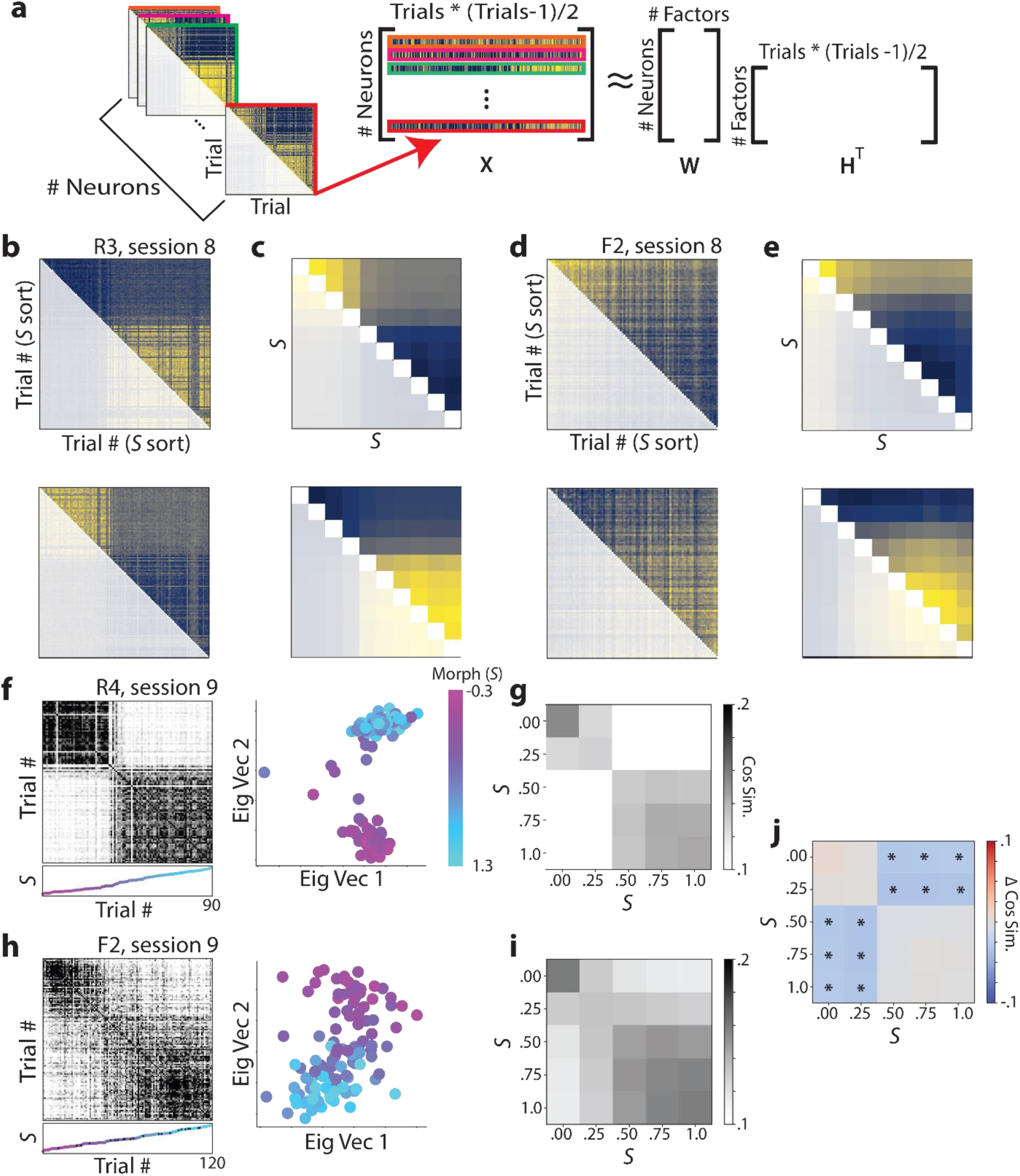
CA1 cells recorded in the Rare Morph condition form two discrete representations for morph values while cells recorded in the Frequent Morph condition form a spectrum of representations for different morph values. **a**, Schematic for how Nonnegative Matrix Factorization (NMF) was performed. For each cell, we flattened the upper triangle of its trial by trial similarity matrix (as seen in Fig 1g & i) to form a vector. These vectors were then stacked to form a *Neurons* × (*Trials* * (*Trials* − 1)/2) matrix, *X*. NMF was then performed on this matrix yielding a factors matrix, *H*^*T*^, and a loadings matrix, *W*. The rows of *H*^*T*^ can be reshaped into “prototypical similarity matrices”. **b**, NMF results for a single Rare Morph session (mouse R3, session 8; 915 cells, 120 trials). Components from the best fitting two component model are shown. Color coding indicates maximum (yellow) and minimum (blue) values. See Extended Data Fig 4 for model comparisons. **c**, NMF results for all Rare Morph sessions and animals (n = 28,409 cells). Trials were binned by morph value and combined across Rare Morph sessions (8-N) and animals. NMF was performed on the resulted set of matrices and the components for the best fitting two component model are shown. **d**, Same as in (**b**) but for an example Frequent Morph session (mouse F2, session 8; 2,127 cells, 120 trials). **e**, Same as in (**c**) for all Frequent Morph sessions (8-N) and animals (n = 19,324 cells). **f**, Left: Population trial × trial cosine similarity matrix for an example Rare Morph session (mouse R4, session 8; 886 cells, 90 trials), color coded for maximum (black) and minimum (white) values. The scatterplot below each similarity matrix indicates the morph value of the trial. Color and height on the vertical axis indicate morph value, with trials sorted in ascending morph value (color bar in the far right side of panel). Black dots indicate trials in which the reward was omitted. Right: The projection of each trial’s activity onto the principal two eigenvectors of the similarity matrix. Each dot indicates one trial, colored by morph value. **g**, Population similarity matrices were binned across morph value and averaged for all Rare Morph Sessions 8-N (mice R1 – R4, n = 21 sessions). **h**, Same as (**f**) for an example Frequent Morph session (mouse F2, session 9; 2149 cells, 120 trials). **i**, Same as (**g**) for all Frequent Morph sessions 8-N (mice F1 – F5, n = 19 sessions). **j**, The difference between the average Rare Morph and Frequent Morph (**g**,**i**) trial by trial similarity matrices. Asterisks indicate significant differences (p < 0.001, permutation test).

Trial by trial population similarity matrices revealed results consistent with those observed at the single cell and population level. For the Rare Morph condition, strong block diagonal structure in the trial by trial similarity matrix suggested clustering of morph values into two representations (example session in Fig 2f, across sessions and mice in Fig 2g; examples from all mice shown in Extended Data Fig 5). While in the Frequent Morph condition, the trial by trial similarity matrices showed a gradual transition between the two extreme morph values (example session in Fig 2h, across sessions and mice in Fig 2i). Indicating that hippocampal remapping patterns differed significantly between the Rare Morph and Frequent Morph condition, the differences in across trial similarity between conditions were significantly away from the block diagonal (Fig 2j, p<.005 two-sided permutation test).

### CA1 remapping approximates optimal estimation of the stimulus

We next asked how similar the CA1 neural representation on a single trial were to the average CA1 neural representation 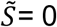 and 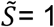 morph values. To quantify this, we defined a Similarity Fraction (SF) between the centroid for the 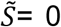 and 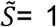 trials. This value is the ratio of the population vector similarity to the 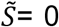 centroid to the sum of similarities to the 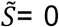 and 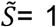 centroids (Fig 3a). Values above .5 indicate trials relatively closer to the 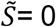 centroid, and values below .5 indicate trials relatively closer to the 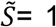 centroid. Plotting SF as a function of the animal’s position along the virtual reality track revealed two stable representations across the entire length of the track for the Rare Morph condition (Fig 3b-c) and a spectrum of representations for the Frequent Morph condition (Fig 3d-e). These results indicate that remapping occurred rapidly upon the start of a new trial and remained stable as the animal traversed the virtual track. This result suggests that we could reasonably consider one SF value for each trial without obfuscating position-dependent accumulation of evidence mechanisms.

**Figure 3:**
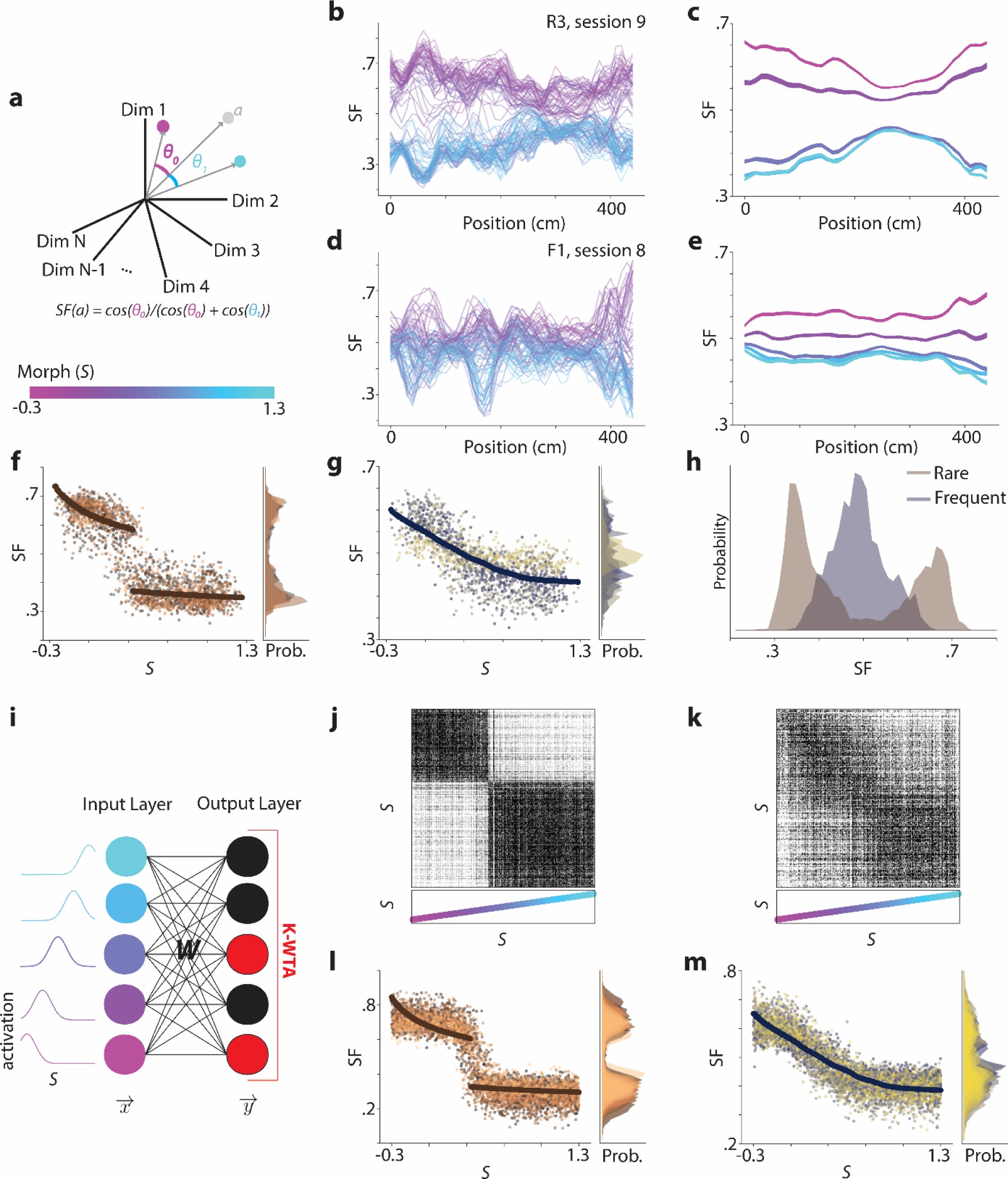
CA1 hippocampal remapping approximates optimal estimation of the stimulus and can be explained by Hebbian learning mechanisms. **a**, *Top:* Schematic for how the Similarity Fraction (SF) is calculated. Each dimension indicates the activity of a single neuron at a single position bin. For panels (**b**-**e**), only one position bin is included in the calculation at a time. The *S* = 0 centroid is calculated by averaging this vector for all *S* = 0 trials (magenta dot). Likewise, the *S* = 1 centroid is calculated by averaging his vector for all *S* = 1 trials (cyan dot). SF for a given trial, *a*, is calculated by dividing the cosine similarity of the vector for *a* to the *S* = 0 centroid by the sum of cosine similarities to either centroid. *Bottom:* Color bar for morph values used in remainder of figure. **b**, SF plotted as a function of the animal’s position on the track for each trial in an example Rare Morph session (mouse R3, session 9; 2046 cells, 100 trials). Each line represents a single trial and color coded indicates morph value (color bar in [**a**]). **c**, Average SF by position plot across Rare Morph sessions (8-N) and animals (R1 – R4; n = 21 sessions). Lines represent means for binned morph values (*S* = 0, .25, .50, .75, 1). Shaded region indicates SEM across sessions. **d**, Same as (**b**) but for an example Frequent Morph session (mouse F1, session 8; 301 cells, 95 trials). **e**, Same as (**c**) but for all Frequent Morph sessions (8-N, animals F1 – F5, n = 19 sessions). **f**, *Left:* Whole-trial SF for each trial was calculated and plotted as a function of morph value for all Rare Morph sessions. Each dot represents a single trial (n = 2,410 trials) and each color represents a different mouse. The bold dots (brown) indicate the cross-validated prediction of SF values from a linear regression of the MAP estimate of the stimulus under the Rare Morph prior (Fig 1e) onto the data. *Right:* Marginal histogram of SF values. Colors correspond to individual mouse. **g**, Same as F but for all Frequent Morph sessions (n = 1,706 trials). The bold dots (navy) indicate the cross-validated prediction of SF values from a linear regression of the MAP estimate of the stimulus under the Frequent Morph prior (Fig 1f). **h**, Histogram of the SF values for all mice in the Rare Morph condition (brown) and Frequent Morph condition (blue). **i**, Schematic of the computational model. The input layer contains neurons that form a basis for representing the stimulus using radial basis functions. Activations are linearly combined via the matrix *W* and thresholding is applied via a K-Winners-Take-All mechanism to achieve the output activation. *W* is updated after each stimulus presentation using Hebbian learning. Model training is performed by drawing trials randomly from either the Rare Morph prior or the Frequent Morph prior. **j**, Trial by trial similarity matrix, as in Fig 2f for experimental data, for an example model trained under the Rare Morph condition. **k**, Same as (**j**) for an example model trained under the Frequent Morph condition. **l**, Whole-trial SF as a function of Morph, as in (**f**), for models trained under the Rare Morph condition. Each color indicates a different random model initialization (n = 50). The bold line is a linear regression of the Rare Morph MAP estimate. **m**, Same as (**k**) but for Frequent Morph trained models (n = 50). The bold line is a linear regression of the Frequent Morph MAP estimate.

Consequently, we calculated a summary SF value for each trial and plotted it as a function of morph value for each session (8-N) and each mouse (“SF-*S* plot”; Fig 3f-g). The distribution of SF values from Rare Morph (n = 2,410 trials) and Frequent Morph (n = 1,706 trials) trials were significantly different (two sample KS test D = .395, p = 4.60 × 10^−128^, Fig 3h). Moreover, SF plots were highly overlapping with the MAP estimate of the stimulus under the two priors (Fig 3f-g, MAP estimate from Fig 1b-c in bold). For the Rare Morph trials, the Rare Morph MAP estimate curve predicted a significant amount of variance in the SF-*S* plot (linear regression, cross-validated R^2^ = .811, p < .001 permutation test). This prediction was significantly better than regressing the Frequent Morph MAP estimate curve or a line to the SF-*S* curve (Extended Data Fig 6a, two-sided Wilcoxon Signed-Rank Tests on magnitude of residuals from cross-validated linear regression; Rare Morph MAP v. Frequent Morph MAP: T = 1.01 × 10^6^, p = 5.73 × 10^−39^, n = 2410, rank-biserial correlation = .244; Rare Morph MAP vs. linear fit: T = 8.37 × 10^5^, p = 1.09 × 10^−72^, n = 2410, rank-biserial correlation = .313). The Rare Morph MAP estimate prediction was not significantly different from fitting a sigmoid to the SF-*S* plot (Extended Data Fig 6a, two-sided Wilcoxon Signed-Rank Test on magnitude of residuals from cross-validated regression, T = 1.42 × 10^6^, p = .360, n = 2410, rank-biserial correlation = .022), but the sigmoid fit failed to capture the clear discontinuity in the data. Similarly, for the Frequent Morph trials, the Frequent Morph MAP estimate curve predicted a significant amount of variance in the SF-*S* plot (linear regression, cross-validated R^2^ = .5336, p < .001 permutation test). This prediction was significantly better than regressing the Rare Morph MAP estimate curve or a line to the SF-*S* curve (Extended Data Fig 6b, two-sided Wilcoxon Signed-Rank Test on magnitude of residuals from cross-validated linear regression, Frequent Morph MAP vs. Rare Morph MAP: T = 6.25 × 10^5^, p = 3.87 × 10^−7^, n = 1706, rank-biserial correlation = .090; Frequent Morph MAP vs linear fit: T = 5.75 × 10^5^, p = 4.81 × 10^−14^, n = 1706, rank-biserial correlation = .155). As above, the Frequent Morph MAP estimate was not significantly different from a sigmoid regression (two-sided Wilcoxon Signed-Rank Test on magnitude residuals from cross-validated regression, T = 6.95 × 10^5^, p = .103, n = 2410, rank-biserial correlation = .041). Together, these results provide evidence that CA1 remapping can be approximated by MAP estimation of the animal’s location in a multidimensional stimulus space in which contextual representations are activated proportional to their probability.

We next sought to implement a network model capable of approximating MAP inference. We reproduced key experimental findings by implementing a model with three main components: 1) stimulus driven input to all cells, 2) Hebbian learning by cells on these inputs, and 3) competition between neurons via K-winners-take-all learning (Fig 3i; See Methods and Extended Data Fig 7 for details and motivation)^14,15^. Using identical model parameters, we simulated learning under the Rare Morph and Frequent Morph conditions by drawing stimuli from the corresponding priors in a one dimensional version of the task. Single model cells showed similar remapping patterns to experimentally observed CA1 cells (Extended Data Fig 7e-f), model-derived trial by trial similarity matrices (Fig 3j-k, Extended Data Fig 7g) bore a clear resemblance to those derived from experimental data (Fig 2f-j) and model-derived SF values across trials (i.e. simulations) were well fit by a MAP estimate curve (Fig 3l-m). This model has the strength of approximating a normative solution to hidden state inference with a plausible learning rule, without explicitly optimizing over an assumed cost function^16,17^.

### Remapping patterns emerge early in training but solidify with experience

We next examined the degree to which the animal’s prior beliefs about states update over time. The hippocampus’s role in memory formation appears critical at the earliest stages of learning^18–21^. However, the previous analyses all considered later (8-N) sessions. To examine remapping earlier in learning, we performed the same similarity analyses on session 3, the first session Rare Morph condition animals experience environments with intermediate morph values. In both the Rare Morph and Frequent Morph condition mice showed variable early session remapping patterns. In the majority of mice (3/4 in the Rare Morph condition, 4/5 in the Frequent Morph condition), the remapping pattern in session 3 was largely similar to the remapping pattern observed in later sessions (example mouse Rare Morph condition Fig 4a-c, m; Frequent Morph condition Fig 4g-i, n). In these animals, a linear regression of the MAP estimate from the matching condition provided a better fit than the MAP estimate curve from the opposite condition. Perhaps due to small numbers of trials, this difference was only significant in animal R2 (two-sided Wilcoxon signed rank test on magnitude of residuals, T = 552, p = 7.53 × 10^−3^, n = 60, rank-biserial correlation = .333) and F3 (T = 2.81 × 10^3^, p = 3.17 × 10^−2^, n = 120, rank-biserial correlation =.150). In contrast, in session 3 of one Rare Morph animal (R1), we observed a gradual transition in the neural representation between S = −0.3 and S = 1.3 morph values, and indeed the Frequent Morph MAP estimate provided a marginally better fit to the data (Fig 4d-f, m). In addition, in session 3 of one Frequent Morph condition mouse (F2), we observed more discrete switches between representations and the Rare Morph MAP estimate provided a marginally better fit than the Frequent Morph MAP estimate (Fig 4j-l). For both of these mice (R1 and F2), the remapping patterns in later sessions (8-N) were akin to those observed in all other animals (i.e. Rare Morph MAP regression fit outperforms Frequent Morph MAP regression fit for R1, two-sided Wilcoxon signed rank test, T = 6.71 × 10^4^, p = 5.42 × 10^−8^, n = 600, rank-biserial correlation = .223; Frequent Morph MAP regression fit outperforms Rare Morph MAP regression fit for F2, T = 3.77 × 10^4^, p = 2.91 × 10^−3^, n = 425, rank-biserial correlation = .101). Together, this indicates that prior-driven remapping patterns are learned early, but strengthen over training.

**Figure 4:**
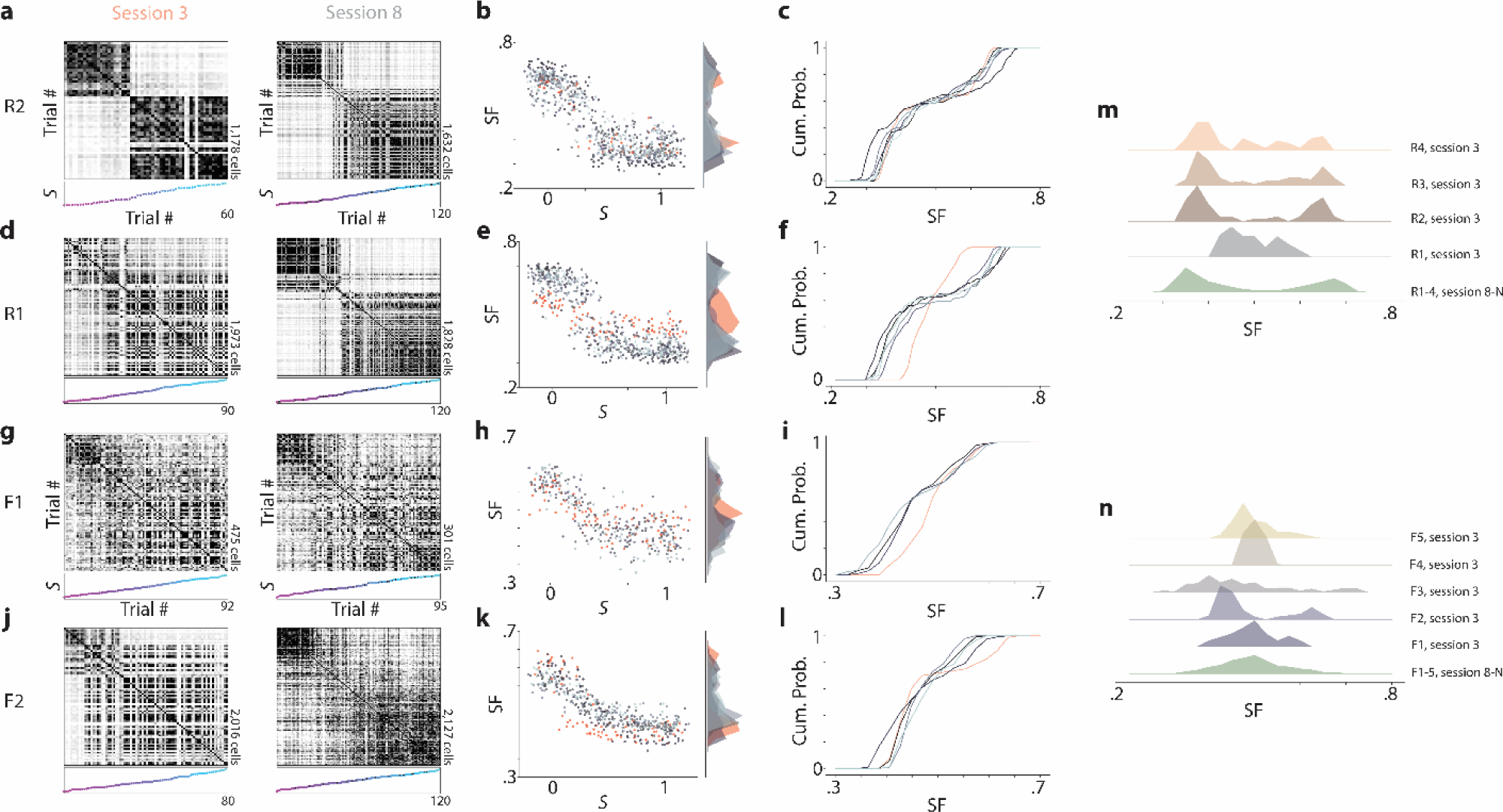
Neural context discrimination is stable across late sessions but forms at variable rates. **a**, *Left:* Trial by trial similarity matrices for session 3 (first session in which the mouse experiences intermediate morph values) from an example mouse in the Rare Morph condition (mouse R2). *Right:* session 8 for the same mouse. **b**, *Left:* SF of each trial plotted as a function of the morph value (*S*). Dots are color coded for session number, with red points indicating data from session 3 and varying shades of grey to blue indicate data from sessions 8-N. *Right:* Marginal histogram of SF values for each session. Color coded as in the left panel. **C)** Cumulative histogram of SF values for the sessions shown in (**b**). **d-f**, Same as (**a-c**) for a mouse in the Rare Morph condition (mouse R1) in which context discrimination dynamics have not stabilized by session 3. **g-i**, Same as (**a-c**) for a mouse in the Frequent Morph condition (mouse F1) in which context discrimination dynamics are stable by session 3. **j-l**, Same as (**a-c**) for a mouse in the Familiar Morph condition (mouse F2) in which the context discrimination dynamics are not stable by session 3. **m**, *Top 4 histograms*: histograms of SF values from session 3 in each Rare Morph mouse (n = 4). *Bottom histogram*: combined histogram of all Rare Morph mice from sessions 8-N for comparison. **n**, *Top 5 histograms*: histograms of SF values from session 3 in each Frequent Morph mouse (n = 5). *Bottom histogram*: combined histogram of all Frequent Morph mice from sessions 8-N for comparison.

Once remapping patterns in CA1 were established, they remained highly robust. In all animals, late sessions displayed stable context discrimination metrics (remaining animal examples in Extended Data Fig 8). This provides further evidence for animals using MAP estimation, as it is the prior and not the amount of accumulated experience in the intermediate trials that correlates with remapping (Extended Data Fig 9). In addition, requiring mice to behaviorally discriminate between morph values greater than .5 versus less than .5, by licking in different spatial locations on the VR track, yielded the same remapping patterns as those observed with the random foraging task (Extended Data Fig 10).

We have shown that the remapping patterns observed in CA1 can be precisely predicted by an animal’s prior experience, with the hippocampus approximating an ideal combination of these prior beliefs with information about the current stimulus. This framework provides a possible unifying explanation for earlier works using morphed 2-dimensional environments, which resulted in conflicting findings regarding remapping in CA1^3,6,10^. Moreover, our computational model demonstrates that our experimental results agree with associative learning mechanisms in the hippocampus, a long-standing proposal for how the hippocampus implements memory^15,22,23^. Broadly, these results complement the recently proposed probabilistic framework of a successor-like representation in the hippocampus and provide a potential mechanism for switching between successor representations for different contexts^11,24^. On the other hand, these results conflict with models that require updates to the path integration coordinate reference frame to evoke remapping, as mice in both Rare and Frequent Morph conditions experienced identical path integration conditions yet showed very different remapping patterns. It remains to be seen whether these dynamics emerge in CA1 or they are inherited from an earlier stage of processing. Our findings build on the emerging idea that the hippocampal-entorhinal circuit represents location not only in physical space but also in abstract stimulus spaces^25,26^. We show that CA1 remapping is a generalization of this phenomenon and provides approximate optimal inference of location in the multidimensional feature space that defines “context”.

## Acknowledgements

This work was supported by funding from the New York Stem Cell Foundation, NIMH MH106475, Office of Naval Research N00141812690, Simons Foundation 542987SPI and the James S McDonnell Foundation to LMG, and a National Science Foundation Graduate Research Fellowship awarded to MHP. We thank A. Diaz for histology and behavioral training assistance. We thank A.H. Williams and S. Ganguli for helpful discussions regarding data analysis and modeling.

## Author Contributions

MHP and LMG conceptualized experiments and analyses. MHP collected and analyzed data and performed computational simulations. MHP and LMG wrote the paper.

The authors declare no competing interests.

## Methods

### Subjects

All procedures were approved by the Institutional Animal Care and Use Committee at Stanford University School of Medicine. Male and female (n = 6 male, 7 female) mice were housed in groups of between one and five same-sex littermates. After surgical implantation, mice were housed in transparent cages with a running wheel and kept on a 12-hour light/dark schedule. All experiments were conducted during the light phase. Mice were between 2 and 5 months at the time of surgery (18.6 – 29.3 grams). Prior to surgery, animals had *ad libitum* access to food and water.

### Calcium indicator expression

Three methods were used to express GCaMP6 in CA1 pyramidal cells. 1) For mice F1-F3, F5, R4, and FD1-FD4 (Extended Data Figure 10 only), hemizygous CaMKIIa-cre mice (Jackson Laboratory stock #005359)^1^ were first anaesthetized by an intra-peritoneal injection of a ketamine/xylazine mixture (8.5 mg/kg). Then, adeno-associated virus containing cre-inducible GCaMP6f under a ubiquitous promoter (AAV1-CAG-FLEX-GCaMP6f-WPRE; Penn Vector Core) was injected into the left hippocampus (500 nL injected at −1.8 mm anterior/posterior [AP], −1.3 mm medial/lateral [ML], 1.4 mm from the dorsal surface [DV]) using a 36 gauge Hamilton syringe (World Precisions Instruments). The needle was left in place for 15 minutes to allow for virus diffusion. The needle was then retracted and the imaging cannula implant was performed (described in the section below). 2) For mice R1-R3, hemizygous CaMKIIa-cre mice were maintained under anesthesia via inhalation of a mixture of oxygen and 0.5 – 2% isofluorane. A retro-orbital injection of AAV-PhP.eB-EF1a-DIO-GCaMP6f (∼2.6 × 10^11 vg per mouse, Stanford Gene Vector and Virus Core) was performed. This retro-orbital injection occurred 20 days prior to the imaging cannula implant (described below). 3) Mouse F4 was a transgenic GCaMP animal (Ai94; CaMKIIa-tTA; CaMKIIa-cre, hemizygous for all alleles; Jackson Laboratory stock #024115) expressing GCaMP6s in all CaMKIIa positive cells.

### Imaging Cannulas and Implant Procedure

We made modifications of previously described procedures for imaging CA1 pyramidal cells^2-4^. Imaging cannulas consisted of a 1.3 mm length stainless steel cannula (3 mm outer diameter, McMaster) glued to a circular cover glass (Warner Instruments, #0 cover glass 3mm diameter; Norland Optics #81 adhesive). Excess glass overhanging the edge of the cannula was shaved off using a diamond tip file.

For the imaging cannula implant procedure, animals were anaesthetized by an intra-peritoneal injection (IP) of a ketamine/xylazine mixture (8.5 mg/kg) and after one hour, were maintained under anesthesia via inhalation of a mixture of oxygen and 0.5 – 1% isoflurane. Before the start of the surgery, animals were also subcutaneously administered 0.08 mg Dexamethasone, 0.2 mg Carprofen, and 0.2 mg Mannitol. A 3 mm diameter craniotomy was performed over the left posterior cortex (centered at −2 mm AP, −1.8 mm ML). The dura was then gently removed and the overlying cortex was aspirated using a blunt aspiration needle under constant irrigation with sterile artificial cerebrospinal fluid (ACSF). Excessive bleeding was controlled using gel foam that had been torn into small pieces and soaked in sterile ACSF. Aspiration ceased when the fibers of the external capsule were clearly visible. Once bleeding had stopped, the imaging cannula was lowered into the craniotomy until the coverglass made light contact with the fibers of the external capsule. In order to make maximal contact with the hippocampus while minimizing distortion of the structure, the cannula was placed at approximated 15 degree roll angle relative to the animal’s skull. The cannula was then held in place with cyanoacrylate adhesive. A thin layer of adhesive was also applied to the exposed skull. A number 11 scalpel was used to score the surface of the skull prior to the craniotomy so that the adhesive had a rougher surface on which to bind. A headplate with a left offset 7 mm diameter beveled window was placed over the secured imaging cannula at a matching 15 degree angle, and cemented in place with Met-a-bond dental acrylic that had been dyed black using India ink.

At the end of the procedure animals were administered 1 mL of saline and .2 mg of Baytril and placed on a warming blanket to recover. Animals were typically active within 20 min and were allowed to recover for several hours before being placed back in their home cage. Mice were monitored for the next several days and given additional Carprofen and Baytril if they showed signs of discomfort or infection. Mice were allowed to recover for at least 10 days before beginning water restriction and VR training.

### Two Photon imaging

To image the calcium activity of neurons, we used a resonant-galvo scanning two photon microscope (Neurolabware). 920 nm light (Coherent Discovery laser) was used for excitation in all cases. Laser power was controlled using a pockels cell (Conoptics). Average power for excitation of AAV1-CAG-GCaMP6f mice and the Ai94;CaMKIIa-tTA;CaMKIIa-cre mouse was 10-40 mW (mouse F1-5, R4, and FD1-3). For AAV-PHP.EB mice the typical laser power was 50-100 mW (mouse R1, R2 and R3). The 1 mm × 1 mm field of view (FOV) (512 × 796 pixels) was collected using unidirectional scanning at 15.46 Hz. Cells were imaged continuously under constant laser power until the animal completed 60-120 trials, the session exceeded 40 min, or the mouse stopped running consistently.

Putative pyramidal cells were identified using the Suite2P software package (https://github.com/MouseLand/suite2p)^5^, and the segmentations were curated by hand to remove ROIs that contained multiple somas, dendrites, or contained cells that did not display a visually obvious transient. This method identified between 98 and 2,149 putative pyramidal neurons per session, depending on the quality of the imaging window implant and expression of the virus. We did not attempt to follow the same cells over multiple sessions; however we attempted to return to roughly the same FOV on each session. For all analyses, we used the extracted “activity rate” obtained by deconvolving the *Δ*F/F with a canonical calcium kernel. We do not interpret this result as a spike rate. Rather, we view it as a method to remove the asymmetric smoothing on the calcium signal induced by the indicator kinetics.

### Virtual Reality (VR) Design

All virtual reality environments were designed and implemented using the Unity game engine (https://unity.com/). Virtual environments were displayed on three 24 inch LCD monitors that surrounded the mouse and were placed at 90 degree angles relative to each other. A dedicated PC was used to control the virtual environments and behavioral data was synchronized with calcium imaging acquisition using TTL pulses sent to the scanning computer on every VR frame. Mice ran on a fixed axis foam cylinder and running activity was monitored using a high precision rotary encoder (Yumo). Separate Arduino Unos were used to monitor the rotary encoder and control the reward delivery system.

### Water Restriction and VR Training

In order to incentivize mice to run, animals water intake was restricted. Water restriction was not implemented until 10-14 days after the imaging cannula implant procedure. Animals were given 0.8 – 1 mL of 5% sugar water each day until they reach ∼85% of their baseline weight and given enough water to maintain this weight.

Mice were handled for 3 days during initial water restriction and watered through a syringe by hand to acclimate them to the experimenter. On the fourth day, we began acclimating animals to head fixation (day 4: ∼30 minutes, day 5: ∼1 hour). After mice showed signs of being comfortable on the treadmill (walking forward and pausing to groom), we began to teach them to receive water from a “lickport”. The lickport consisted of a feeding tube (Kent Scientific) connected to a gravity fed water line with an in-line solenoid valve (Cole Palmer). The solenoid valve was controlled using a transistor circuit and an Arduino Uno. A wire was soldered to the feeding tube and capacitance of the feeding tube was sensed using a simple RC circuit and the Arduino capacitive sensing library. The metal headplate holder was grounded to the same capacitive-sensing circuit to improve signal to noise, and the capacitive sensor was calibrated to detect single licks. The water delivery system was calibrated to deliver ∼4 *μ*L of liquid per drop.

After mice were comfortable on the ball, we trained them to progressively run further distances on a VR training track in order to receive sugar water rewards. The training track was 450 cm long with black and white checkered walls. A pair of movable towers indicated the next reward location. At the beginning of training, this set of towers were placed 30 cm from the start of the track. If the mouse licked within 25 cm of the towers, it would receive a liquid reward. If the animal passed by the towers without licking, it would receive an automatic reward. After the reward was dispensed the towers would move forward. If the mouse covered the distance from the start of the track (or the previous reward) to the current reward in under 20 seconds, the inter-reward distance would increase by 10 cm. If it took the animal longer than 30 seconds to cover the distance from the previous reward, the inter-reward distance would decrease by 10 cm. The minimum reward distance was set to 30 cm and the maximal reward distance was 450 cm. Once animals consistently ran 450 cm to get a reward within 20 seconds, the automatic reward was removed and mice had to lick within 25 cm of the reward towers in order to receive the reward. After the animals consistently requested rewards with licking, we began Rare Morph or Frequent morph training described below. Training (from first head fixation to the beginning Rare Morph or Frequent Morph protocols) took 2 – 4 weeks.

### Morphed Environments

For the Rare and Familiar Morph condition experiments, trained mice ran down 450 cm tracks in order to receive sugar water rewards. These rewards were placed at a random location between 250 and 400 cm down the track. The reward location was indicated by a small white box with a blue star. Animals received a reward if they licked within 25 cm of the box. Once the animals were well trained they often ran consistently down the first half of the track and begin licking as they approached the reward (Extended Data Fig 1). They typically stopped to consume the reward and ran at a consistent speed to the end of the track. Well-trained mice ran until they received around 0.8 – 1 mL of liquid (∼200 rewards). In order to increase the number of trials in a session, rewards were omitted on 20% of trials for some sessions. Reward omissions or missed rewards are shown in the results when necessary.

The visual stimuli for the VR track were chosen so that the extremes of the stimulus distributions could be gradually and convincingly morphed together. The tracks did not change in length or the location of salient landmarks. The aspects of the stimulus that did change were, i) the frequency and orientation of the sine waves on the wall (low frequency, oriented sine waves into high frequency nearly vertical sine waves), ii) the color of the first two towers (green to blue), and iii) the color of the background of the visual scene (light grey to dark grey). The morph value, *S*, for each trial is the coefficient of an affine combination between two extreme values of the stimulus (*Sf*_0_ + (1 − *S*)*f*_1_). On every trial, a shared morph value, 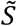, was chosen for all features from one of five values (0, .25, .5, .75, and 1) and a jitter was applied independently to the wall, tower color, and background colors. The jitters were uniform random values from -.1 to .1. The total morph value is given by the sum of the shared morph value and all of the jitters (shared morph + wall jitter + background jitter + tower jitter). The resulting range of *S* values is between −0.3 and 1.3, though values close to −0.3 and 1.3 will be rare.

### Rare Morph Condition

Mice in the Rare Morph condition experienced only trials with a shared morph value of 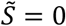 or 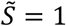, with the exception of a subset of imaging sessions. Only one session was run per day. For the first two sessions, the animals experienced trials with randomly interleaved 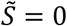 or 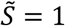 trials. On the third session, before imaging, the animals experienced 30-50 ‘warm-up’ trials of with randomly 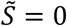 or 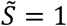 trials. During imaging (60-120 trials, depending on the running speed of the animal), 50% of the trials were 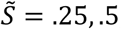, or. 75 trials. The remaining 50% were 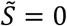 or 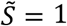 trials. After the imaging session, animals continued to run 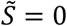 and 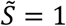 trials until they received the rest of their water for the day. For the next four sessions (4-7), the animals again only experienced 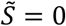 and 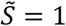 trials. For session 8 and all subsequent sessions (8-N) the protocol used in the third session was repeated.

### Frequent Morph Condition

Mice in the Frequent Morph condition experienced randomly interleaved 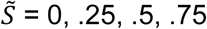 or 1.0 trials with equal probability on every session. On imaging sessions, as in the Rare Morph condition, animals experienced 30-50 ‘warm-up’ trials before imaging and continued trials after imaging until they received the rest of their water for the day.

### Log-Morph Calculation

According to Weber-Fechner’s Law, the percept of a stimulus is proportional to the log of the stimulus intensity. Therefore, despite the fact that the stimulus values are linearly interpolated, the mouse likely does not perceive the morph in a linear fashion. To correct for this, we calculated a new morph value based on the log-scale of the stimuli. We first calculated the log of each possible feature value in its native range and then normalized so that this value was between −0.3 and 1.3, *log S*.

### Maximum A Posteriori Morph (MAP) Inference

MAP inference is the process of taking the maximum value from the posterior distribution using Bayes’ formula. The posterior distribution, in this case, describes the probability of the stimulus, *S*, given an observation, *Ŝ*. We can calculate the posterior using *P*(*S*|*Ŝ*) ∝ *P*(*Ŝ*|*S*)*P*(*S*). *P*(*S*) is the prior distribution and describes the frequency of previously seen stimuli. The Rare Morph and Frequent Morph conditions were designed to vary this distribution and are described below. *P*(*Ŝ*|*S*) is the likelihood function which describes observation noise. In all cases, we assume that this function is a Gaussian centered at *S* in the log-morph scale with a variance of .3, *P*(*S*) ∝ *N*(*S*, .3). We then compute the MAP estimate as *P*(*S*|*Ŝ*). In Fig 1e-f, we plot this value as a function of *Ŝ*.

For the Rare Morph condition, *P*(*S*) ∝ *G*(−.1, .1) + *G*(.9,1.1) + *c*. Where *G*(*a, b*) *= U*(*a, b*) * *U*(*a, b*) * *U*(*a, b*). The * operator denotes a convolution, *U*(*a, b*) is a uniform distribution over the interval *[a, b]*, and *c* is constant. The convolutions come from the fact that the total morph value is the sum of three features with the same distribution and the property that the distribution of the sum of random variables is proportional to the convolution of the distributions. We add a constant, *c*, to account for intermediate trials seen in session 3 and to prevent the probability of seeing an intermediate stimulus from being exactly 0. For the Frequent Morph condition, *P*(*S*) ∝ *G*(−.1, .1) + *G*(.15, .35) + *G*(.4, .6) + *G*(.65, .85) + *G*(.9,1.1) + *c.*

Both priors were then smoothed with a small Gaussian kernel (variance = .1) and converted to the log-morph scale where the MAP estimation was performed.

### Place cell identification and plotting

Place cells were identified separately in 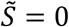 and 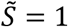 trials using a previously published spatial information (SI)^6^ metric 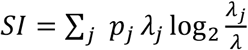. Where *λj* is the average activity rate of a cell in position bin *j, λ* is the position-averaged activity rate of the cell, and *p*_*j*_ is the fractional occupancy of bin *j*. The track was divided into 10 cm bins, giving a total of 45 bins. To prevent cells that were active on only a small number of trials from having spuriously high spatial information, we performed a bootstrapping procedure for estimating *λ*_*j*_ and *λ*. On each iteration of the bootstrap (30 total iterations), 67% of the trials were chosen with replacement and the SI was calculated from the cellular activity in these trials. The median SI across bootstrapping iterations was taken as the final SI value.

To determine the significance of the SI value for a given cell, we created a null distribution for each cell independently using a shuffling procedure. On each shuffling iteration, we circularly permuted the cell’s time series relative to the position trace within each trial and repeated the bootstrapping procedure to determine SI for the shuffled data. Shuffling was performed 1000 times for each cell, and only cells that exceeded all 99% of permutations were determined to be significant “place cells”.

Place cell sequences (Fig 1h & j) were found separately in the 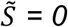 and 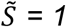 trials using split-halves cross-validation. The average firing rate maps from a randomly selected half of the trials were used to identify the position of peak activity. Cells were sorted by this position and then the activity on the other half of the trials was plotted. This gives a visual impression of both the reliability of the place cells within the 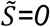 and 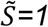 morphs and the extent to which these sequences are retained across the intermediate morph values.

For visualization, single trial activity rate maps (Fig 1g & i) were smoothed with a 20 cm (2 spatial bin) Gaussian kernel.

### Single Cell Trial × Trial Similarity Matrices

For each cell, *i*, we stack smoothed single trial activity rate maps (10 cm/1 spatial bin width Gaussian kernel) to form a matrix, *A*_*i*_ ∈ *R*^*T,J*^. *T* is the number of trials and *J* is the number position bins. Each row was then divided by its *l*_2_norm, yielding a new matrix *Ã*_*i*_. The single cell trial by trial cosine similarity matrix is then given by *C*_*i*_ *= ÃÃ*_*i*_ ^*T*^

### Population Trial × Trial Similarity Matrices

For a single session, we horizontally concatenated all single cell trials by positions matrices, *A*_*i*_, to form the fat matrix *A =* [*A*_1_|*A*_2_| … |*A*_*N*_], where *N* is the number of neurons recorded in that session. To calculate the population trial × trial cosine similarity matrix, we again divide the rows of *A* by their *l*_2_norm to give the matrix *Ã*. As above, the population cosine similarity matrix is then given by *C = ÃÃ*^*T*^. In order to average across sessions, the rows and columns of *C* were binned by morph value.

### Non-negative Matrix Factorization

For each session, we applied a previously described cross-validated version of non-negative matrix factorization (NMF)^7^ to the set of matrices *C*_1_, *C*_2_, …, *C*_*N*_ described above to identify dominant remapping patterns seen across the population. To perform NMF, we flatten the upper triangle of each matrix *C*_*i*_ giving a row vector 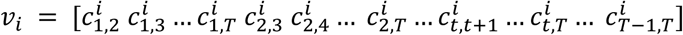 and stack them to form matrix 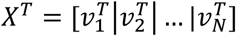, so *X* ∈ *R*^*N,T*(*T*−1)/2^. The goal of NMF is to find matrices, a skinny matrix *W* and a fat matrix *H*^*T*^, that minimize 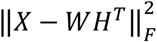. We can control the number of factors we get out of the model by specifying the rank of *W* and *H*. In short, to perform cross-validation, we applied a random binary masking matrix *M* during training and test the model on the left out data. The optimization then becomes 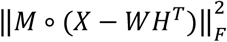, where 70% of the entries in *M* are 1 and the remainder are 0. The ○ operator indicates the elementwise product. Performance of the model was then tested by examining 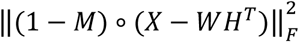. Cross-validation allows us to find the best number of factors to include in our model, as test error should plateau or get worse when the rank of the model exceeds the “true” dimensionality. We performed three fold cross-validation for all of our models.

### Similarity Fraction

We define a Similarity Fraction (SF) to quantify the relative distance of a single trial representation to either the average 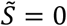 or 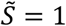 representation. For whole-trial SF, on each trial, *t*, we take the *t*^*th*^ row of *A, α*_*t*_. We calculate the cosine similarity between *α*_*t*_ and the average 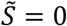 population representation, 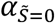 given by 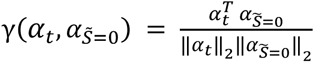. Similarly we calculate the cosine similarity between *α*_*t*_ and the average 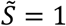 population representation, 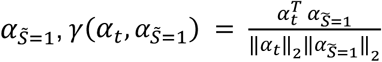. If *t* is a 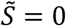 or 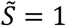 trial, it is omitted from the centroid calculation. SF is then given by 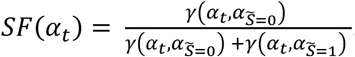. To calculate SF as a function of position, we simply considered only the columns of *A* that correspond to that position bin.

For fitting the MAP estimates onto the SF values, we performed robust linear regression (Huber loss function) where the MAP estimate at a given trial’s morph value was the feature and we predict the actual SF value for that trial. These regressions were compared to a standard linear fit to the SF plots, and a Huber loss sigmoid fit to the data. For all model comparisons, we examined Leave-One-Trial-Out residuals.

### Hebbian Learning Model of Data

Our goal for this model was to write down the simplest network with plausible components that would replicate our main findings and approximate MAP inference. We posited that this would be possible without any explicit optimization and would only require three main components: 1) stimulus driven input neurons, 2) Hebbian plasticity on input weights by output neurons and, 3) competition between output neurons. For this model we considered a one-dimensional, position independent version of the task.

We considered a two layer neural network with a set of stimulus-driven input neurons, 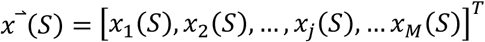, a set of output neurons,*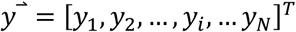*, and a connectivity matrix *W*, where *W*_*ij*_ is the weight from input neuron *j* to output neuron *i*. Input neurons have radial basis function tuning for the stimulus, 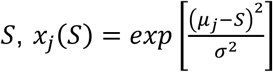, where *μ*_*j*_ is the center of the radial basis function for neuron *j*. These basis functions were chosen to tile the stimulus axis across the population, 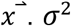 is the width of the radial basis function and is a fixed value across all input neurons. For Fig3i-m & Extended Data Fig 7d-h, M = 100, N = 100, and *σ*^2^ *=* .15.

We accomplished competition between output neurons using a K-Winners-Take-All approach. On any given stimulus presentation *y*_*i*_ *= max*{*c*_*y*_*KWTA*(*z*(*S*)) ○ *z*(*S*) + *σ*_*y*_, 0} where 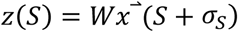, *KWTA*(·) is a vector-valued function that applies the K-Winners-Take-All threshold and outputs a binary vector choosing the K winners, *c*_*y*_ is a constant, and *σ*_*y*_ is additive noise. *σ*_*s*_ is a stimulus noise term. For Fig 3i-m & Extended Data Fig 7d-h, K=40 (approximately the same fraction of place cells observed in the real data), *c*_*y*_ *=* .01, *σ*_*y*_∼2 * *N*(0,1), *σ*_*S*_∼.05 * *N*(0,1)

Weights were largely updated according a basic Hebbian learning rule, *ΔW*_*ij*_ *= ηx*_*j*_*y*_*i*_, where *η* is a constant. However, we also required that weights cannot be negative, that weights cannot go beyond a max value, *W*_*max*_, and that weights decay at some constant rate, *τ*. This yielded the following update equation *W*_*ij*_ := *min*{*max*{*W*_*ij*_ + *ΔW*_*ij*_ − *τ*, 0}, *W*_*max*_}. For Fig 3i-m & Extended Data Fig 7d-h, *η =* .15, *σ*_*w*_∼.1 * *N*(0,1), *τ =* .005, *W*_*max*_ *=* 10.

For each instantiation of the model, we initialize *W* with random small weights (*W*_*ij*_∼⋃(0,.1)) and training stimuli are chosen according to either the Rare Morph prior or the Frequent Morph prior (n=1,000 training stimuli). After training, *W* is frozen and we test the models response to the full range of stimuli.

## Extended Data Figures

**Extended Data Figure 1.**
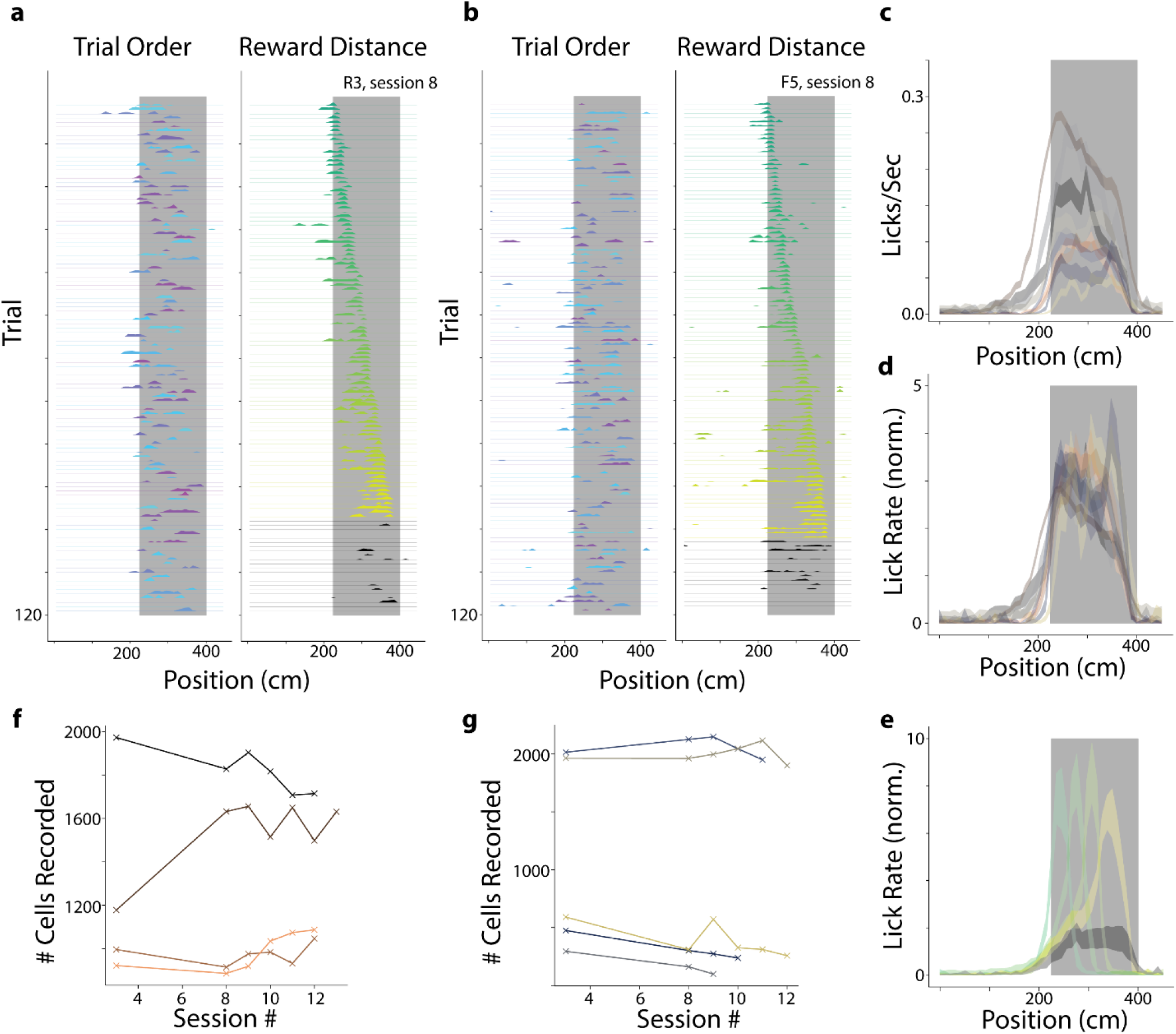
Licking behavior and number of cells recorded per session. **a**, Single trial lick rate as a function of position is shown for an example Frequent Morph session (F5, session 8; n= 120 trials). *Left-*Each row indicates the smoothed lick rate across positions for a single trial. The color of the rows indicate the morph value (colormap in Fig 1a). The grey shaded region indicates possible reward locations. Trials are shown in the order in which they occurred during the experiment. *Right-*Trials are sorted by the location of the reward cue. The color code also indicates increasing reward distance from the start of the track (green to yellow). Black trials are those in which the reward cue was omitted. **b**, Same as (**a**) for an example Rare Morph session (R3, session 8; n=120 trials). **c**, Across session mean lick rate (licks/sec) as a function of position (see [**f**-**g**] for number of sessions and color code) is plotted as a separated line for each mouse (R1-R4 & F1-F5; mean ± SEM). The color scheme for each mouse is the same as in the rest of the manuscript. Vertical shaded region indicates possible reward location as in (**a-b**). **d**, The same data as (**c)** is normalized by the animal’s overall mean lick rate. **e**, Normalized mean lick rates were combined across Rare and Frequent Morph animals. Trials were then binned by reward location (50 cm bins) and plotted as a function of position (across animal mean ± SEM). Color code is the same as (**a-b**). **f**, The number of cells identified per session is plotted for each Rare Morph animal individually. Each mouse shown as a different color. **g**, Same as (**f**) for all Frequent Morph animals.

**Extended Data Figure 2.**
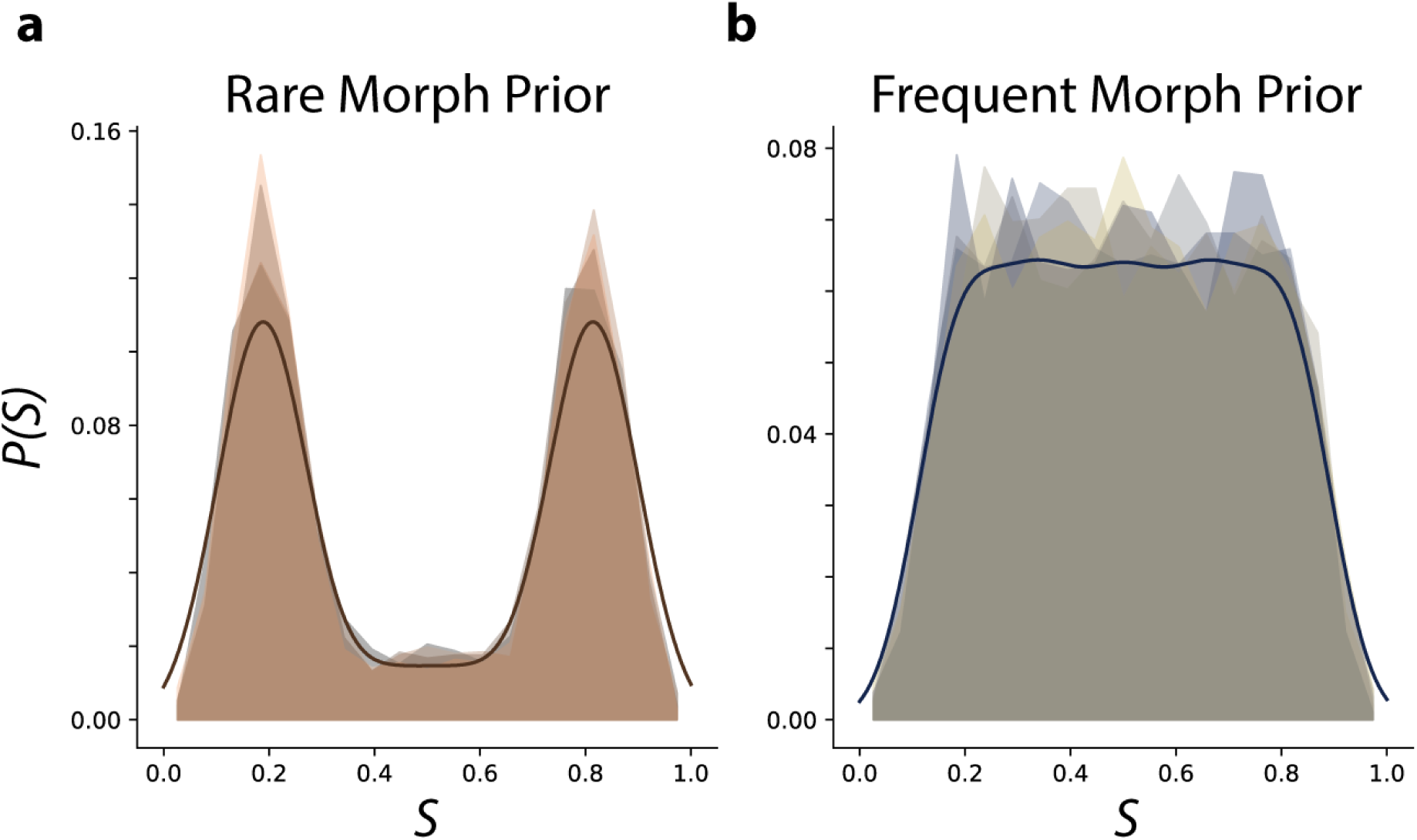
Empirical and idealized priors over morph values **a**, The probability of a morph value occurring across all sessions (1-N) is shown for each Rare Morph animal as a normalized histogram. Animal color scheme as in Extended Data Fig 1f. The idealized prior (see methods) used for MAP estimates is shown in bold. **b**, Same as (**a**) for all Frequent Morph animals. Animal color scheme as in Extended Data Fig 1g.

**Extended Data Figure 3.**
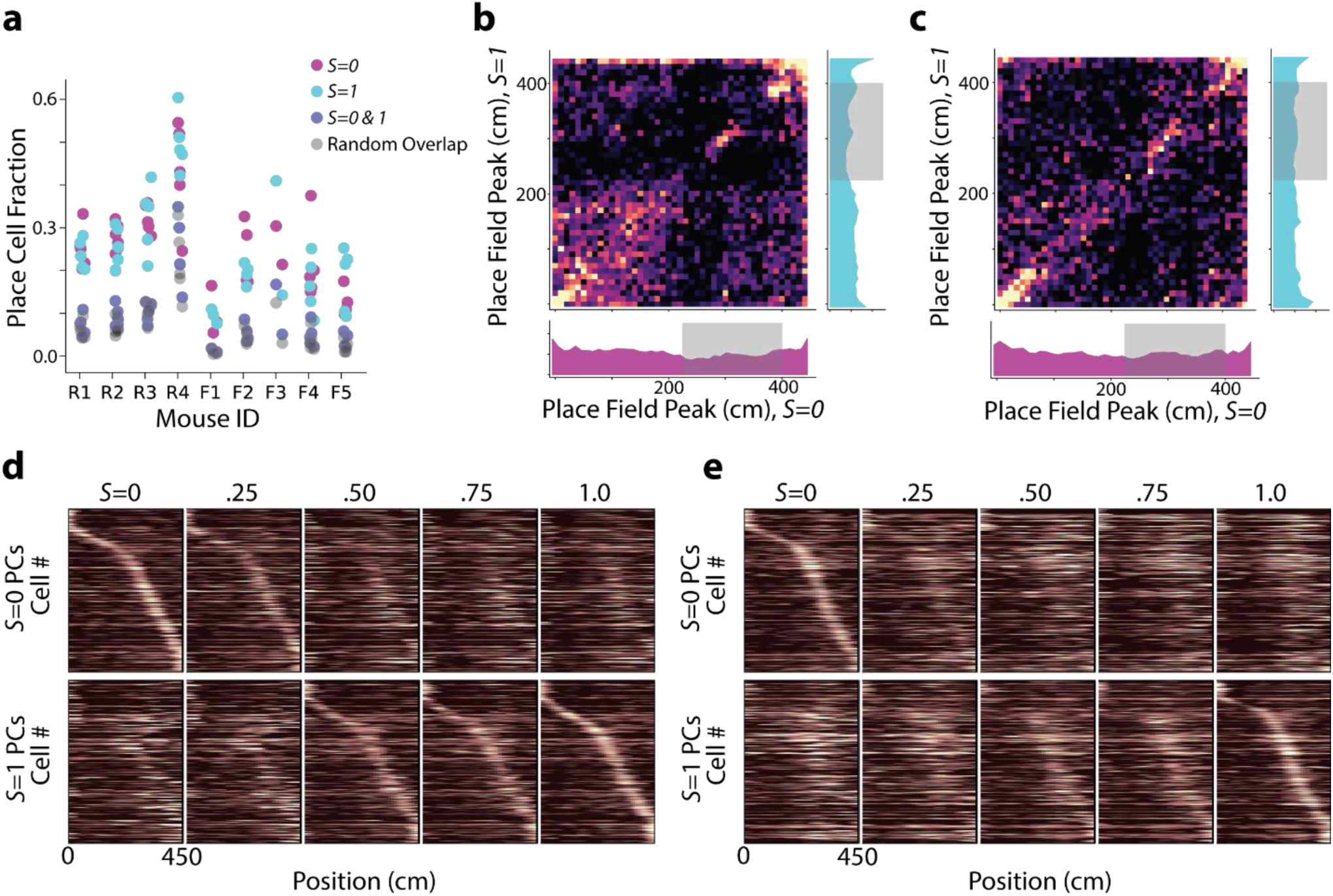
Analysis of place cells with place fields in both *S* = 0 and *S* = 1 environments. **a**, Fraction of cells classified as place cells in the S = 0 trials (magenta), S = 1 trials (cyan), and both S = 0 and S = 1 trials (intersection of previous two sets of cells; navy). The number of cells expected to be classified as place cells in both environments if cells were chosen randomly with replacement is also shown (grey). **b**, For cells that were classified as place cells in both S = 0 and S = 1 trials (n = 3,239 cells) in Rare Morph animals, we plot the location of peak activity for S = 0 trials against the location of peak activity for S = 1 trials. Remapping appears largely random except for populations coding for the beginning and end of the track and reward locations. Marginal histograms are shown for occupancy of position bins in the *S* = 0 trials (bottom) and *S* = 1 trials (right). Shaded regions indicate possible reward locations. **c**, Same as (**b**) for all Frequent Morph animals (n = 1,040 cells). **d**, Same as Fig 1h but only cells that had place fields in both S=0 and S=1 trials are plotted (n = 3,239 cells). **e**, Same as Fig 1j but only cells that had place fields in both S = 0 and S = 1 trials are plotted (n = 1,040 cells).

**Extended Data Figure 4.**
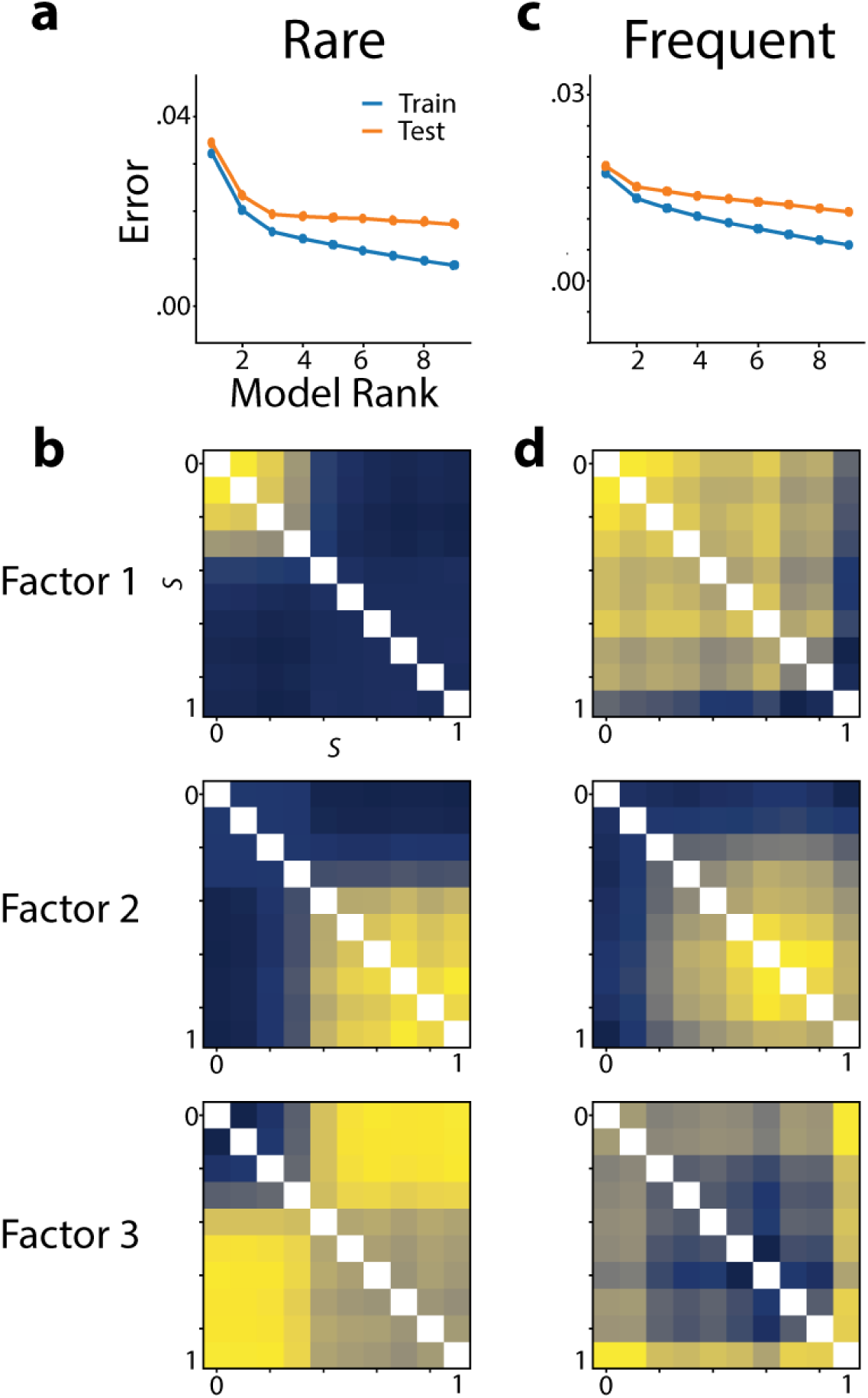
Higher rank NMF models and error analysis (related to Fig 2a-e). **a**, Training and test reconstruction error for NMF models binned by morph value and combined across all Rare Morph sessions (related to Fig 2c, see methods for details, 3 folds of cross-validation). **b**, Trial × trial similarity matrix factors from the best rank 3 NMF model for Rare Morph cells. This yields a qualitatively similar result as Fig 2c with an additional factor for the level of off-diagonal similarity in each cell. **c**, Same as (**a**) for all Frequent Morph sessions (related to Fig 2e). **d.** Same as (**b**) for best rank 3 NMF model for Frequent Morph cells. The first two factors show a gradual transition from one representation to the other. Factor 3 appears to isolate off diagonal similarity as in the Rare Morph rank 3 model.

**Extended Data Figure 5.**
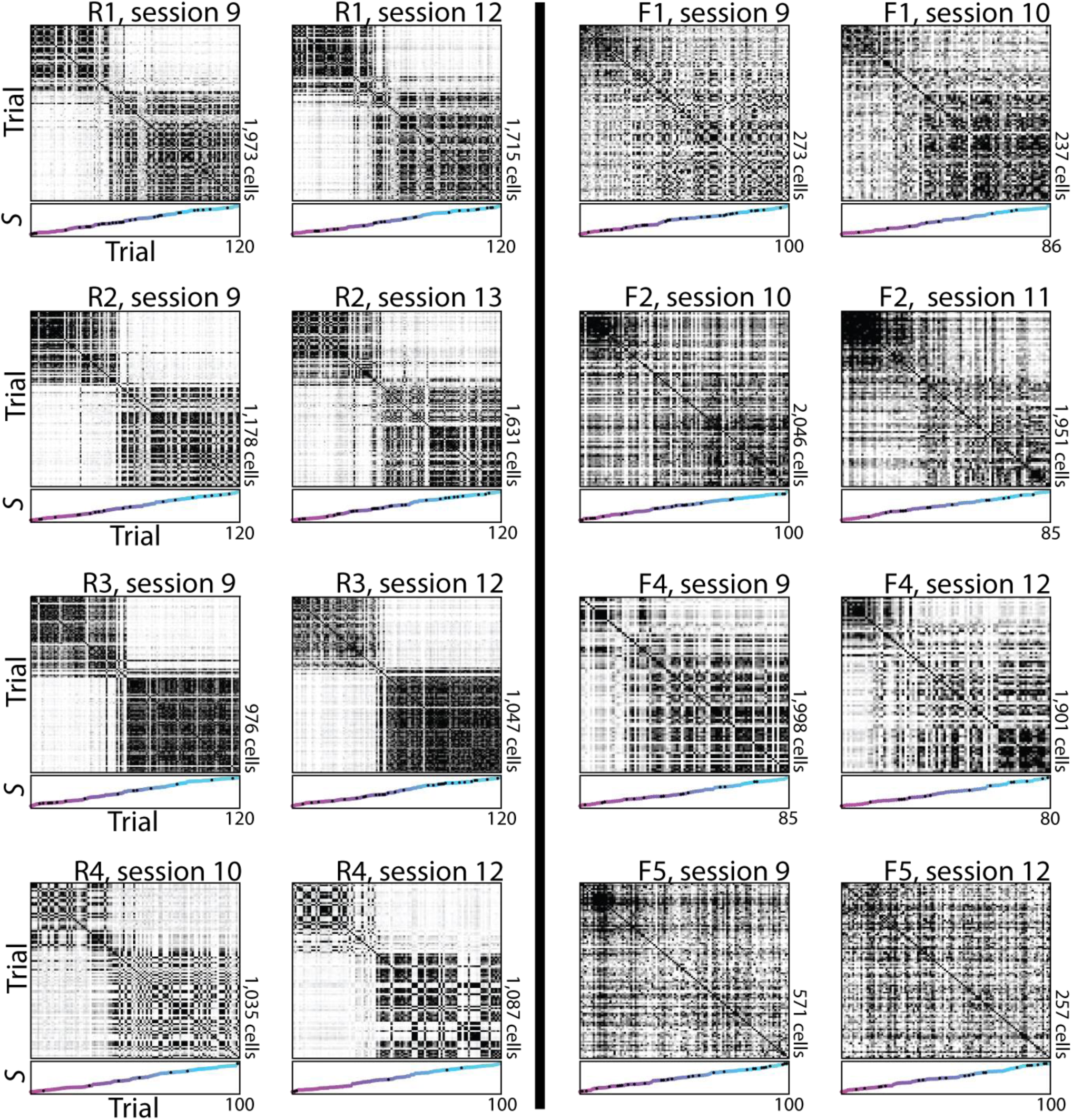
Example session trial by trial similarity matrices. For each animal, (*Left-*Rare Morph, *Right-*Frequent Morph) we show two example session trial by trial similarity matrices, as in Fig 2f & h. Each matrix is sorted by ascending morph value. We show the earliest session not shown elsewhere in the manuscript and the last imaging session.

**Extended Data Figure 6.**
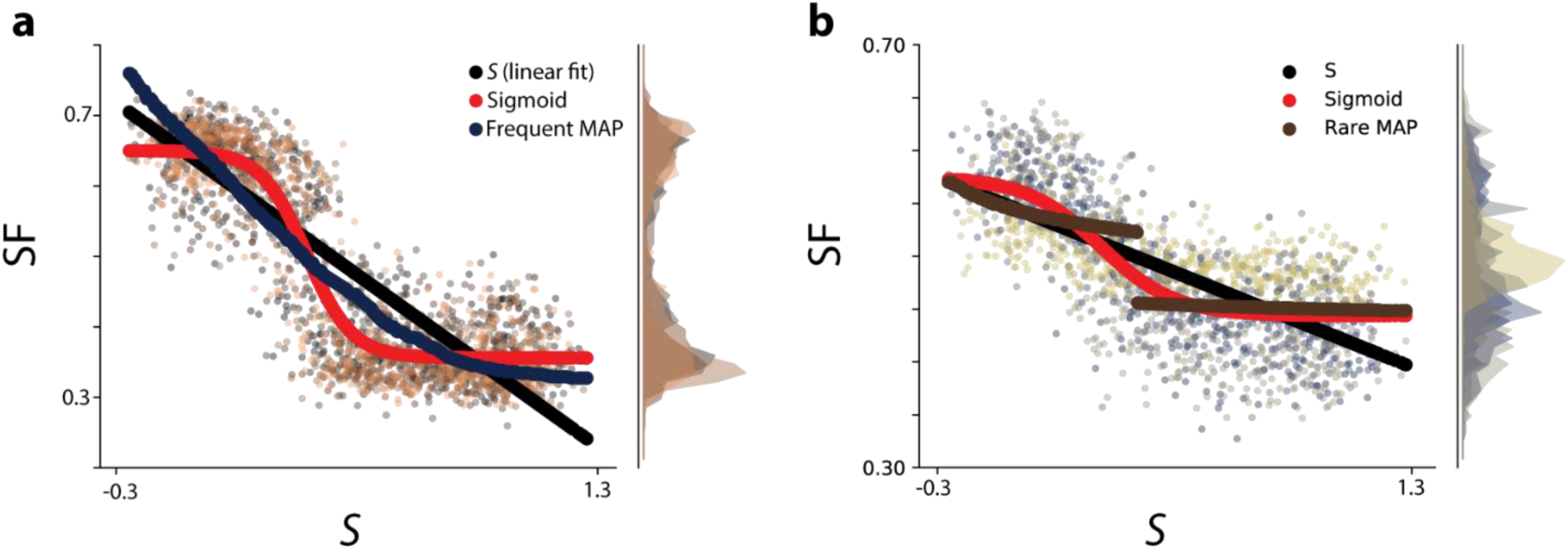
Comparison of regressions onto Similarity Fraction (SF) vs Morph data (related to Fig 3f-g). **a**, SF is plotted as a function of morph, *S*, for all Rare Morph sessions (8-N, colors indicate animal) as in Fig 3f. The cross-validated prediction from a linear regression (black), a sigmoid fit (red), and a linear regression of the Frequent Morph MAP estimate (navy) are shown. **b**, Same as (**a**) for all Frequent Morph sessions. Instead of the Frequent Morph MAP estimate, the Rare Morph MAP estimate regression (brown) is shown.

**Extended Data Figure 7.**
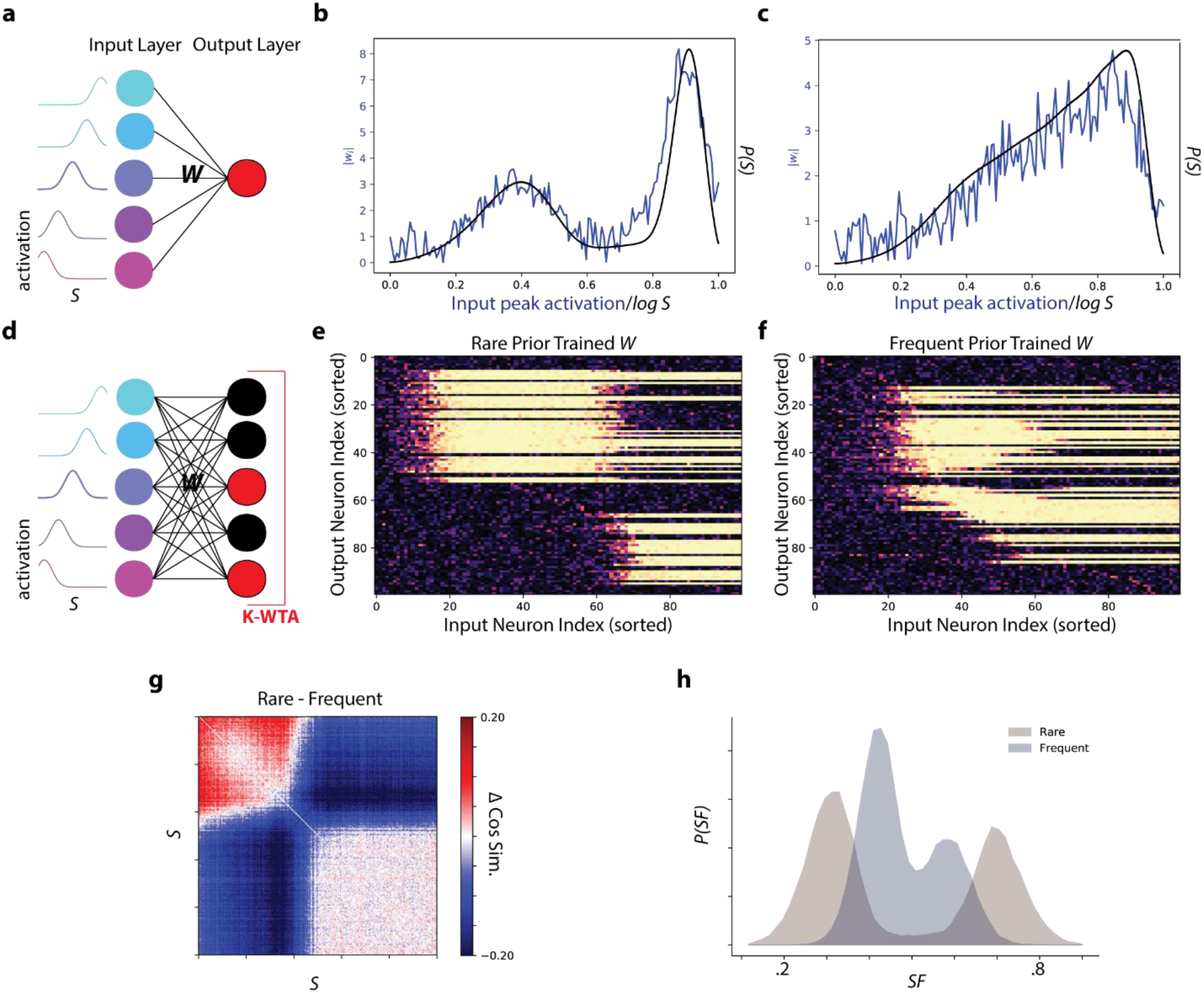
Motivation for network model and additional characterization of model behavior. **a**, Schematic of single cell version of model. A single output cell receives morph-tuned inputs. All other details of the model are the same as the multi-output cell version. In addition, since there is only one output cell, there is no “Winner-Take-All” mechanism. **b**, The single cell model can learn the prior over morph values from which the stimuli are drawn. For a model trained with the Rare Morph prior, the weights (*w*_*i*_) onto the output cell are plotted as a function of the peak selectivity of the input cells (blue). For comparison, the prior distribution from which the stimuli were drawn is plotted as well (black). **c**, Same as (**b**) for a Frequent Morph trained single cell model. **d**, Schematic of the multiple output cell model (replica of Fig 3I). The single cell model can learn the prior over presented stimuli, but the output is not particularly useful for inferring the stimulus. By adding competition between the output cells, we can distribute this prior across cells and readout the posterior from the activity of the population. Adding the “Winner-Take-All” mechanism forces the network to represent the most likely stimulus, and thus approximate MAP inference. **e**, The weight matrix, *W*, of a Rare Morph trained model. The input neuron index, *i*, is indicated by the columns (sorted by peak morph selectivity), and the output neuron index, *j*, is indicated by the rows of the matrix (sorted by magnitude of peak input weight). The heat indicates the value of the weight. Weights are largely selective for either lower morph values or higher morph values. Output cells inherit this behavior and remap discretely like the cells from Rare Morph animals. **f**, Same as (**e**) for a Frequent Morph trained model. Similar to the real data, cells show more gradual changes in selectivity across morph values. **g**, Difference in average trial by trial similarity matrices for Rare Morph trained models and Frequent Morph trained models (similar to Fig 2j). **h**, Histogram of SF values for the Rare Morph trained (brown) and Frequent Morph trained (navy) models (similar to Fig 3h).

**Extended Data Figure 8.**
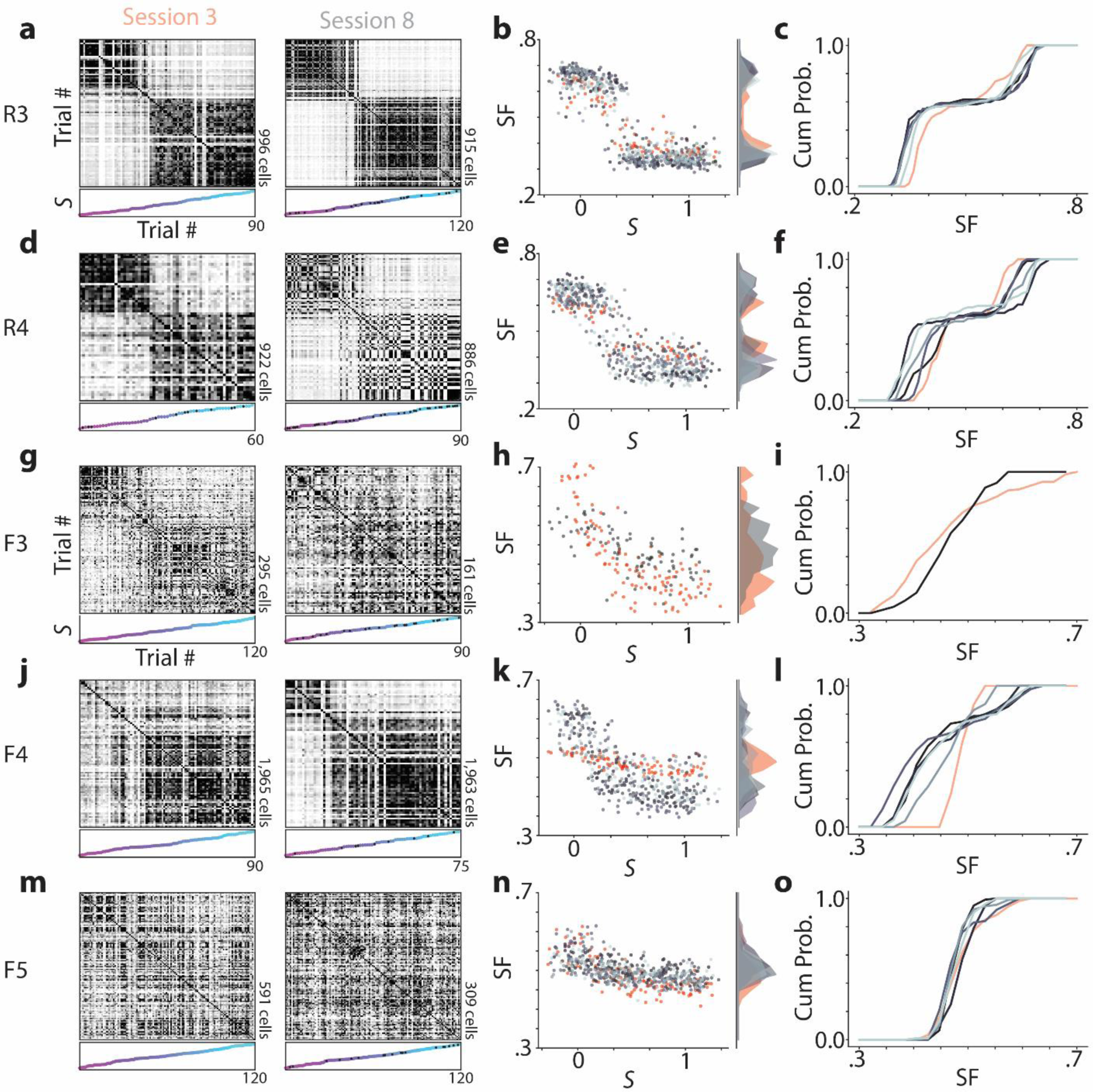
Additional examples of early vs late context discrimination (related to Figure 4). **a**, *Left:* Trial by trial similarity matrices for session 3 (first session in which the mouse experiences intermediate morph values) from an example mouse in the Rare Morph condition (mouse R3). *Right:* session 8 for the same mouse. **b**, *Left:* SF of each trial plotted as a function of the morph value (*S*). Dots are color coded for session number, with red points indicating data from session 3 and varying shades of grey to blue indicate data from sessions 8-N. *Right:* Marginal histogram of SF values for each session. Color coded as in the left panel. **C)** Cumulative histogram of SF values for the sessions shown in (**b**; same as Fig 4a-c). **e-f**, Same as Fig 4a-c for animal R4. **g-i**, Same as Fig 4g-i for animal F3. **j-l** Same as Fig 4g-i for animal F4. **m-o** Same as Fig 4g-i for animal F5.

**Extended Data Figure 9.**
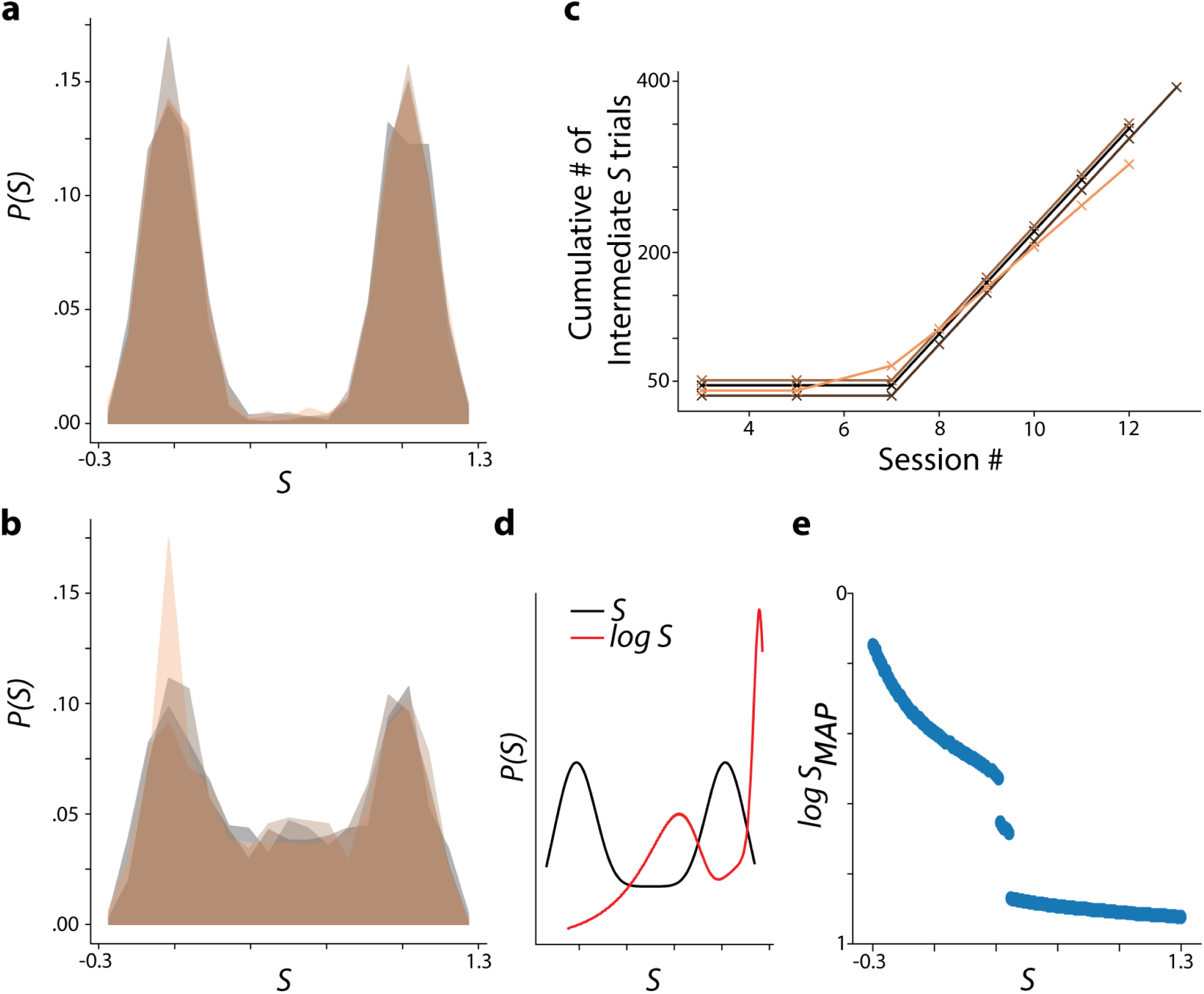
Rare Morph MAP estimation remains qualitatively stable despite cumulative experience with intermediate morph trials. **a**, Rare Morph prior for early sessions (1-7). Colors indicate individual animals as in Extended Data Fig 1f. **b**, Rare Morph prior late sessions only (8-N). **c**, Cumulative number of intermediate morph trials (S =.25 to .75) over sessions for each Rare Morph animal. A small vertical jitter was added to make all curves visible. **d**, Idealized Rare Morph prior for late sessions only (8-N, black) and log correction for this prior (red; similar to Fig 1e). **e**, Log-corrected MAP estimate of the morph value, *log SMAP*, as a function of the true morph value, *S*.

**Extended Data Figure 10.**
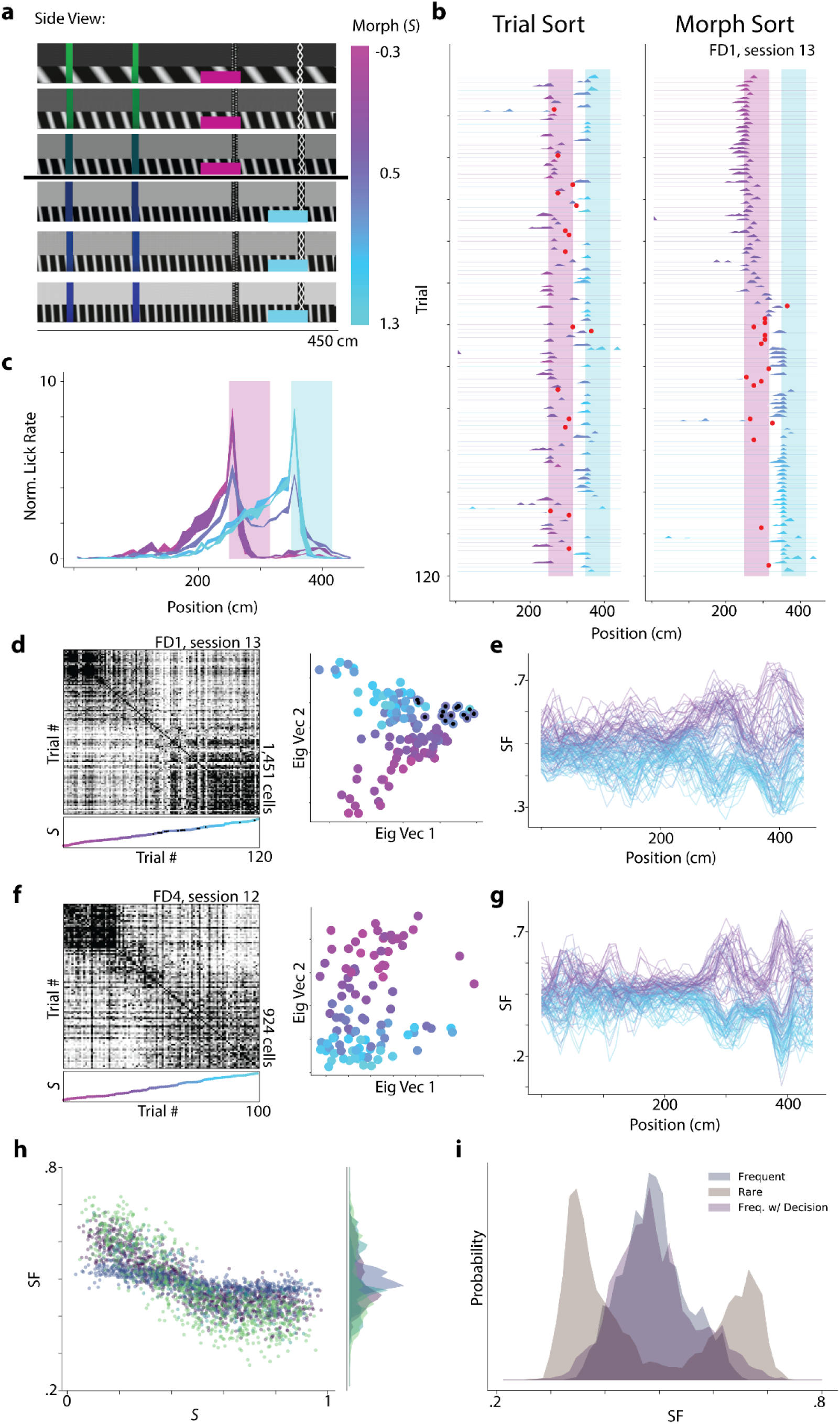
Requiring Frequent Morph animals to behaviorally categorize morph values does not change neural context discrimination. **a**, Stimulus design, as in Fig 1a, for Frequent Morph trained animals that had to behaviorally categorize morph values (Frequent Morph with Decision, n = 4 animals, FD1 – FD4). A side view of a subset of VR tracks with different morph values (*S*) are shown. Vertical location indicates approximate morph value for the track shown (color bar to the right of tracks). For trials where *S ≤* .5, animals had to lick within a 65 cm region surrounding the third tower (polka dot tower, magenta highlighted region). For trials where S > .5, animals had to lick within a 65 cm region surrounding the fourth tower (hatched pattern tower, cyan highlighted region). For one animal (FD1), we added punishments for licking in the incorrect reward zone after session 3. If the animal licked in the incorrect reward zone, it was instantly teleported to a dark hallway for 10 seconds before being able to begin the next trial. **b**, *Left:* Lick rate as a function of position for each trial as in Extended Data Fig 1a-b for an example session (FD1, session 13, n = 120 trials). Red dots indicate error trials in which the mouse licked in the incorrect reward zone and was given a timeout. Color indicates morph value of the trial. Shaded regions indicate reward zones as in (**a**). *Right*: Trials are sorted by increasing morph value. Most errors are made near *S* = .5, and appear to be due to inability to withhold licking for the last few cm of the reward zone instead of licking in anticipation of the wrong reward zone. **c**, For mice that did not receive timeouts for incorrect licks (n = 3, FD2-4), we plot the across mouse average normalized lick rate (normalization as in Extended Data Fig 1) as a function of position (across mouse mean ± SEM) for binned morph values (*S* = 0, .25, .5, .75, 1). Shaded regions indicate reward zones. We can see that anticipatory licking behavior is largely categorical with the exception of trials the reward transition (*S* = .5). **d.** *Left:* Trial by trial cosine similarity matrix sorted by increasing morph value for an example session from the mouse that experience timeouts (F1, session 13; 120 trials, 1451 cells). *Right*: Projection of single trials onto the principal two eigenvectors of the similarity matrix. Color indicates morph value. Black dots indicate error trials. Error trials are shifted in the horizontal axis, but this is due to zero-padding the population vector for spatial bins after the teleportation. **e**, Similarity Fraction (SF) as a function of position for each trial for the same example session as in (**d**). SF has a larger magnitude near reward zones. This may be due to context invariant reward coding cells that shift their position of firing across the contexts (see Extended Data Fig 3). **f**, Same as (**d**) for an example session from an animal that did not receive timeouts for incorrect licks (F4, session 12; 100 trials, 924 cells). **g**, Same as (**e**) for the example session shown in (**f**). **h**, *Left:* SF as a function of morph value, *S*, for all late sessions (8-N) from animals FD1-FD4 as in Fig 3f-g (n = 2,762 trials). Each color indicates a different mouse. *Right:* Marginal histogram of SF values for each mouse. **i**, Histogram of SF values for the three different experimental cohorts (Rare Morph-brown, Frequent Morph-navy, Frequent Morph with Decision-purple).

